# Identification of allele-specific KIV-2 repeats and impact on Lp(a) measurements for cardiovascular disease risk

**DOI:** 10.1101/2023.04.24.538128

**Authors:** S. Behera, J. R. Belyeu, X. Chen, L. F. Paulin, N.Q.H. Nguyen, E. Newman, M. Mahmoud, V. K. Menon, Q. Qi, P. Joshi, S. Marcovina, M. Rossi, E. Roller, J. Han, V. Onuchic, C. L. Avery, C.M. Ballantyne, C. J. Rodriguez, R. C. Kaplan, D. M. Muzny, G. A. Metcalf, R. Gibbs, B. Yu, E. Boerwinkle, M. A. Eberle, F. J. Sedlazeck

## Abstract

The abundance of Lp(a) protein holds significant implications for the risk of cardiovascular disease (CVD), which is directly impacted by the copy number (CN) of KIV-2, a 5.5 kbp sub-region. KIV-2 is highly polymorphic in the population and accurate analysis is challenging. In this study, we present the DRAGEN KIV-2 CN caller, which utilizes short reads. Data across 166 WGS show that the caller has high accuracy, compared to optical mapping and can further phase ∼50% of the samples. We compared KIV-2 CN numbers to 24 previously postulated KIV-2 relevant SNVs, revealing that many are ineffective predictors of KIV-2 copy number. Population studies, including USA-based cohorts, showed distinct KIV-2 CN, distributions for European-, African-, and Hispanic-American populations and further underscored the limitations of SNV predictors. We demonstrate that the CN estimates correlate significantly with the available Lp(a) protein levels and that phasing is highly important.

## Introduction

Cardiovascular disease (CVD) is the leading cause of mortality in the world (16.00%) and is on the rise, killing 365,914 people in the United States in 2017 alone ^1^. CVD kills one person every 36 seconds in the United States ^2^. The majority of CVD risk is genetically predetermined but can be abated with proper treatment and lifestyle changes ^3–5^. Quantifying CVD disease risk is challenging and understanding the genomic complexity of adult diseases such as CVD is the next frontier in genomics. Many genomic variants have been associated with CVD risk across dozens of genes, comprising both rare Mendelian effects and more common complex interactions ^6^. Deciphering rare variants and decoding these complex interactions are both challenges that rely on high-accuracy variant calling, consistent phenotyping, and, often, large cohort sizes ^7^. For best results, these studies must consider all types of human genomic variation, including single nucleotide variants (SNVs), structural variants (SVs), and copy number variants (CNs), with both coding and non-coding variations. While difficult to detect and assess, SVs and CNs are the largest source of genetic diversity ^8^ and have been shown to impact human diseases in many cases ^9^.

Numerous studies, including meta-analyses of cohort data, have shown the association between high lipoprotein(a) (Lp(a)) concentrations and CVD ^10,11^. Unlike other traits impacting CVD risk, Lp(a) concentrations are primarily driven by the *LPA* gene ^12^. *LPA* contains multiple Kringle domains, including a hypervariable ∼5,500 bp Kringle IV-type 2 (KIV-2) region, which occurs in tandem from fewer than six copies (the copy number present in Genome Reference Consortium Human Build 38) to at least 50 times ^10^. Furthermore, the KIV-2 region includes two exons that are translated and thus directly impact Lp(a) isoforms. KIV-2 CN allele length has been shown to be inversely associated with increased CVD risk in some populations ^13–15^. While other factors play a role, such as SNVs in non-KIV-2 regions of *LPA*, it has been shown that higher KIV-2 copy number generally associates with decreased Lp(a) protein abundance and CVD risk, and vice versa, except at unusually short allele lengths ^16,17^. Heritability of Lp(a) blood plasma concentrations have also been reported from 70% - 90% ^18–20^, indicating the generational consistency of *LPA* allelic impact on Lp(a) abundance, and the likelihood of Lp(a)-driven CVD risk being similarly heritable. This demonstrates the importance of directly assessing KIV-2 CN states for analysis of CVD risk.

KIV-2 also exhibits >95% length polymorphism in most populations ^10^. Analysis using the T2T-CHM13 reference, which contains 22 copies of the KIV-2 repeat, found a median of 36 copies of KIV-2 in a cohort of 268 genomes, with a standard deviation of approximately 8.5 copies ^21^. This degree of repeat polymorphism also associates with variable Lp(a) levels in both European and African-descended cohorts ^22^. Individuals of African descent have 2-3 fold higher Lp(a) concentration than most individuals from European or Asian populations ^10^.

Despite the importance of Lp(a), it remains unknown why there is such extensive Lp(a) variation. The exact variants that underlie Lp(a) concentration are also not well-characterized. This can be explained in part by the fact that the KIV-2 repeats can account for 70.00% or more of the coding sequence of *LPA* and are inaccessible by common sequencing and genotyping techniques ^23^. It is thus unknown how many functional variants are hidden inside the KIV-2 region, and their potential contribution to the observed diversity of Lp(a) is difficult to predict.

The complexity of the KIV-2 repeat has led to the identification of several marker SNVs in *LPA* but outside the repeat region ^10,24,25^. These markers have been reported to lie in strong linkage disequilibrium with certain KIV-2 CN states. For example, rs10455872 + rs3798220 are often used as marker SNVs and explain around 36.00% of Lp(a) variation in Europeans ^10^, leading to their use in commercial kits to determine CVD risk. However, these SNVs have shown no association in Japanese (only other population studies) and Hispanic populations ^10,26,27^. The marker SNVs are generally absent in autochthonous Africans, low frequency in African-Americans and Europeans, but are in high frequency in Asian and mixed Americans (Mexicans, Colombians, Puerto Ricans, Peruvians) ^25^. While some SNV seems to be functional within their population they often show no effect or are totally absent in other populations^10^. Thus, many SNV markers are only in strong linkage disequilibrium (LD) within a certain population, demonstrating a lack of cross-population portability. This makes them unreliable predictors of Lp(a) levels in most genomes.

To obtain deeper insights into *LPA* KIV-2 region and its consequences on Lp(a) we developed a copy number (CN) calling method, the DRAGEN LPA caller, using Illumina WGS. Comparing the DRAGEN LPA caller against optical mapping, we show that this caller can accurately determine the total number of copies of the KIV-2 repeat from standard whole-genome sequencing. This caller not only enables scaling and routine assessment of KIV-2 on widely available Illumina data but further is able to phase CN of KIV-2. This provides a more detailed insight into the haplotype diversity of KIV-2 per sample and leads to interesting consequences on the protein level. We used the DRAGEN LPA caller to investigate KIV-2 CN across 1000 Genomes Project (1KGP) samples, revealing important insights into a large and diverse cohort. This highlighted that the haplotype diversity of Europeans for *LPA* is quite high compared to other populations. Using this data set we further investigated the relationship of 24 proposed marker SNVs and the actual copy number of KIV-2 across multiple populations. We then utilized the DRAGEN LPA caller to study *LPA* across 3,006 Americans, including 1,001 Hispanics. For 2,375 individuals we had detailed Lp(a) plasma measurements and medical records. This analysis revealed important insights into *LPA* and the related CVD risk for minorities. We leveraged the phasing of KIV-2 CN values to improve the representations, tested association, and here report insights for the impact of KIV-2 repeats on Lp(a) directly across multiple populations. Thus this work provides a methodology to ease the assessment of KIV-2 that improves our understanding of the KIV-2 *LPA* region and its consequences on the protein level Lp(a) and thus CVD risk.

## Results

The complex repeat structure of KIV-2, with a repeat unit of over 5.5 kbp and a median of 36 copies per genome^21^, makes the investigation of *LPA* challenging. Mapping in this region is challenging even for long reads (see **Figure 1A**). Many reads in each size category map non-uniquely to the KIV-2 region (indicated by reads shown in white), leading to low reliability of WGS data in this region.

**Figure 1:**
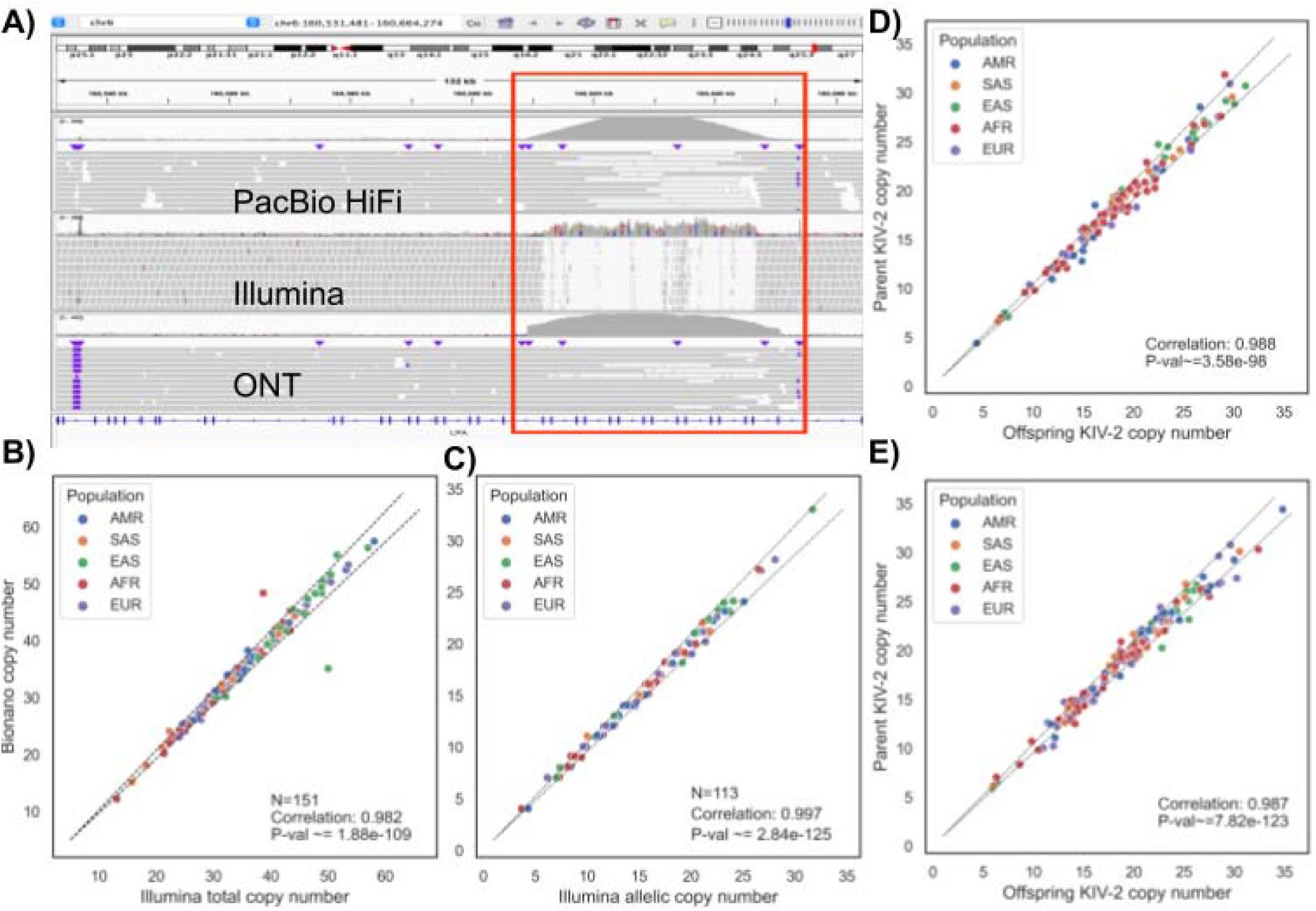
Overview of KIV-2 LPA performance: **A)** Genome wide alignment of short Illumina reads and long PacBio HiFi and ONT reads to LPA gene. We observed many non-unique mappings (i.e. mapping quality MQ=0, shown in white) in KIV-2. The GRCh38 representation of KIV-2 contains six copies of the 5.5 kbp repeat, challenging even for longer reads. This is shown by many white (MQ=0) PacBio reads and to a lesser extent for Oxford Nanopore Technologies (ONT) reads. The fraction of non-unique mapped reads increased with shorter reads, which often hindered the direct assessment of KIV-2 repeat copy number. **B)** Total KIV-2 copy number compared between calls made by the DRAGEN LPA caller or using optical mapping technology, for cases where optical mapping spanned both genomic alleles. Dashed lines indicate error margins of 5% from optical mapping copy number. **C)** Allelic KIV-2 copy number compared where the per-allele copy number was available from the DRAGEN LPA caller and from optical mapping assemblies. Dashed lines indicate error margins of 5% from optical mapping copy number. **D)** For 60 trios where both offspring and parent KIV-2 allele lengths are reported, an allele combination is chosen which minimizes the total difference between each of the two offspring alleles and one from each parent as the most likely allele origin. Each pair, consisting of one offspring allele and one associated parent allele, is shown for a total of 120 allele pairs. Dashed lines indicate error margins of 5% from expected. **E)** For 153 duos where both offspring KIV-2 allele lengths and those from one parent are reported, a most likely allele combination for one offspring allele and one parental allele is chosen as in C. Dashed lines indicate error margins of 5% from expected

We designed the DRAGEN LPA caller to enable direct measurement of KIV-2 copy numbers and thus improve our understanding of *LPA* diversity and impact across a large collection of samples. This tool uses depth of coverage and specialized normalization to determine the copy number of the KIV-2 repeat unit. It then utilizes predetermined SNVs within KIV-2 (**Supplementary Figure S1**), when heterozygously present in a genome, to phase copy numbers per haplotype. A detailed description of the KIV-2 caller can be found in the methods section. The KIV-2 caller is also included in the DRAGEN variant calling software (v4.2). Using a ∼30x Illumina-sequenced whole genome, it requires approximately four minutes to predict the KIV-2 CNs.

### Illumina DRAGEN KIV-2 calls compared against orthogonal data

We compared our DRAGEN LPA caller against Bionano optical mapping results from the Human Genome Structural Variant Consortium ^28,29^, including a total of 12 samples, and an additional set of 154 genomes optically mapped for this comparison. This led to a total set of 166 optically mapped genomes for comparison. While the optical mapping data provides the best high-throughput assay available to assess larger regions of the genome (read size n50 of ∼290 kbp), the *LPA* region still proved challenging even for optical mapping. This challenge was driven in part by the length of the repeat array and in part because the KIV-2 region does not include the nuclease recognition sites used in Bionano optical mapping, leading to incidents of failed contig assembly (see **Supplementary Figure S2**). Optical mapping data succeeded in reporting both alleles in 151 cases, while in 15 cases a single allele was reported. In a small number of additional samples, neither allele could be spanned; these samples were dropped from the comparison analysis. Despite these difficulties, optical mapping provided a large set of high-confidence comparison calls for benchmarking DRAGEN LPA caller results (**Supplementary Table 1)**.

**Figure 1B** shows a significant correlation between high-confidence total copy number calls from the DRAGEN LPA caller and Bionano optical mapping (correlation: 0.982, p-value ∼=1.88e-109). The significant correlation held across all populations tested here. The median difference in total copy number between the two approaches was approximately 0.47 copies, (1.2% of total allele length). The caller was able to identify the allele-specific copy number in 47.06% (1,507/3,202) samples from the 1KGP cohort (**see Supplementary Tables 2 and 3**). Allele-specific copy number call comparisons, in **Figure 1C**, also yielded high correlations (correlation: 0.997, p-value ∼=2.84e-125). An additional set of 96 samples with low-confidence KIV-2 CN calls from Bionano are reported in **Supplementary Figure S2** and **Supplementary Table 1**.

### Diversity of KIV-2 repeats across the human population

We next expanded our study to assess the different KIV-2 copy numbers in different populations. Studies are often focused on European populations or other single populations only so studying KIV-2 across a broad spectrum of the human population provides new insights. For this we assessed the KIV-2 CN estimates among all 3,202 samples (**see Supplementary Tables 2 and 4**) of the 1KGP dataset that includes five different ancestral populations: African (AFR), American (AMR), European (EUR), East Asian (EAS) and South Asian (SAS).

Where possible, we examined the consistency of allele-specific CN calls in the 1KGP trios. Our KIV-2 caller identified phased (i.e. allele-specific) CNs in all samples across 60 trios (**Supplementary Table 5**). This allowed us to compare inherited proband KIV-2 allele lengths to their parental alleles. **Figure 1D** shows the results given one allele from each parent, selected based on the smallest size difference. Among 120 alleles in the trio comparison, the size difference between the observed and inherited allele is within ±5% in 93 cases (77.5% of alleles, correlation coefficient 0.988, significant at □=0.05).

In another 153 trios, the DRAGEN LPA caller identified allele-specific CNs in the proband and one parent. This allowed us to compare one proband allele with the closest-matching parental allele, the most probable allele origin (**Figure 1E**). The size difference between the observed and inherited allele is within ±5% in 115 cases (75.2% of alleles, correlation coefficient 0.987, significant at □=0.05). This translates to a CN difference of <1 given a median haplotype CN of 15. This supports previous studies showing that the KIV-2 CN is indeed highly inherited ^25,30^, including across different populations.

Given the human population catalog of KIV-2 CN across the 3,202 individuals, (**Figure 2A**) we next investigated the distributions across the different populations. First, we studied the distribution among 633 European population samples, which belonged to five subgroups (CEU, FIN, GBR, IBS, and TSI). The majority (∼70.00%) of the population have KIV-2 CN in the range of 30-45 with a mean CN estimate of 37.5 and a standard deviation of 6.86. We used a two-sample Kolmogorov-Smirnov test to compare the CN distribution of EUR samples with other samples. The EUR and SAS populations have almost identical distribution with p-value = 0.047. The distribution was slightly different when compared to AFR and AMR (as shown in Empirical Cumulative Distribution Function (ECDF) plot, **Supplementary Figure S3**) with p-values 1.567e-08 and 7.336e-06 respectively. However, we observed a larger difference when compared to EAS samples (p-value < 2.2e-16). The ECDF plot (**Supplementary Figure S4**) of the haplotype difference i.e., absolute difference of allele-specific CN, showed the distribution of AMR is slightly different from other populations. We further studied the distribution among five different subgroups within the European population and observed that the distributions were very similar among these subgroups (see **Supplementary Figure S5**).

**Figure 2.**
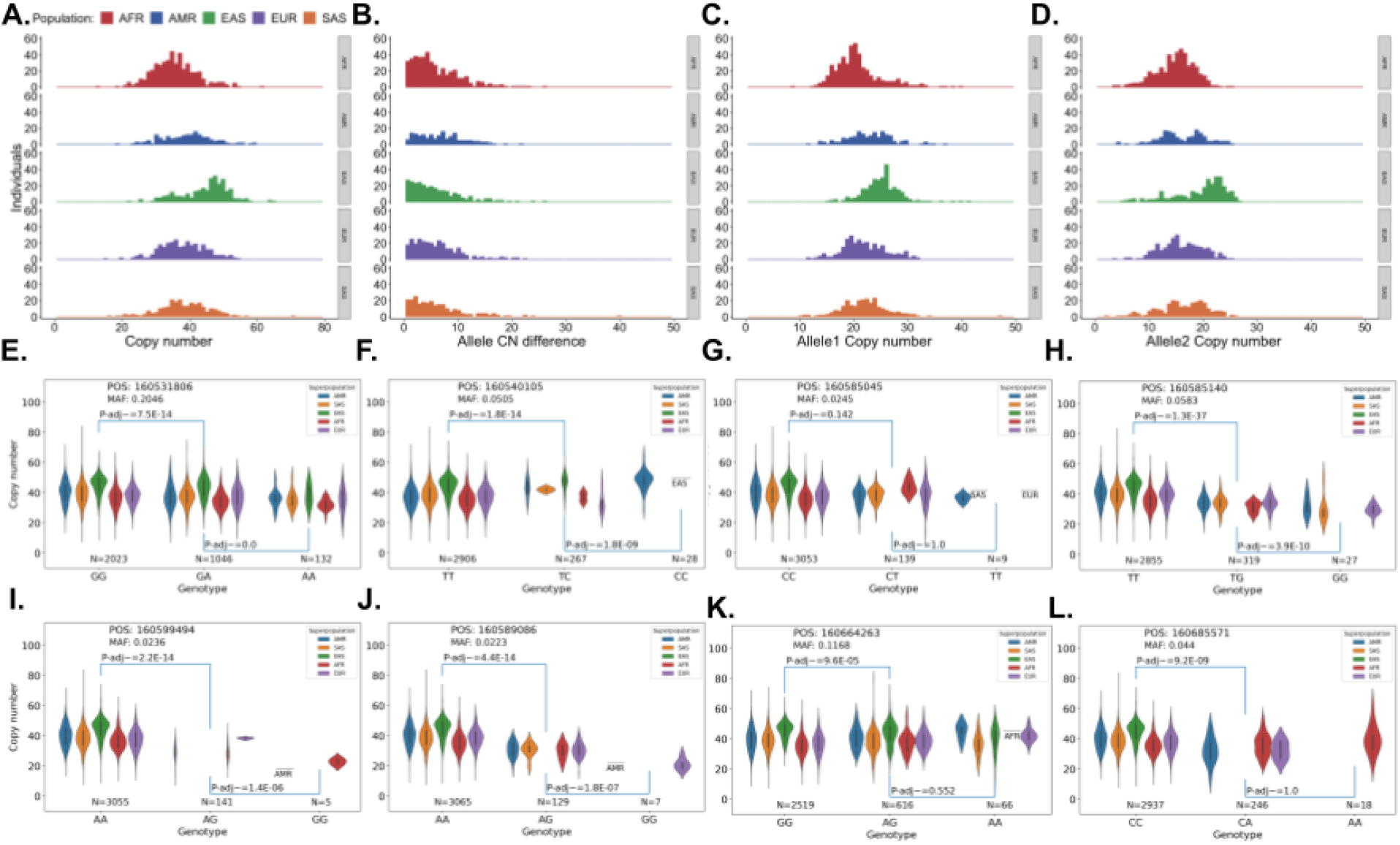
DRAGEN LPA caller analysis across 1KGP cohort. **A)** Distribution of LPA KIV-2 repeats among samples from 1KGP. **B)** Histogram of absolute value of allele length difference between Allele1 and Allele2 in 1KGP samples. The DRAGEN LPA caller could resolve haplotype copy number in about 50% of samples. **C)** The distribution of Allele1 (longest allele in sample) copy numbers among haplotype-resolved 1KGP samples. **D)** The distribution of Allele2 copy numbers (shortest allele in sample). **E-L)** Comparison of SNV markers with CNV states of KIV-2. Samples from 1KGP were genotyped for 12 SNVs which have been reported as associated with different Lp(a) levels or copy number states. KIV-2 copy number was compared between groups of samples homozygous for the major (more common) or minor (less common) alleles of these SNVs, to test for association between SNV status and KIV-2 copy number. P-values were obtained by Kolmogorov-Smirnov tests and corrected for multiple testing by the Bonferroni method.

We expanded our study on the KIV-2 CN distribution among other non-European populations such as Africans, Americans, Asians (South and East). For the AFR dataset, the mean CN was 35.70 with a standard deviation of 6.68. The distribution of CNs among the African population was observed to be different from other populations with several peaks in the range of 30 to 40. The distribution among seven different subgroups within the African population is shown in **Supplementary Figure S6**. The distributions of AMR and EAS are significantly (p-value < 2.2e-16) different compared to AFR. The higher Kolmogorov-Smirnov statistic i.e. D=0.55552 of EAS samples confirms that distribution of EAS samples is different from AFR. For AMR samples, the Kolmogorov-Smirnov statistic when compared to AFR was 0.26431. We have the lowest number (490) of samples for the AMR group and we observed that the AMR (shown in **Figure 2B**) population has a flatter distribution as compared to others. The mean CN estimate for AMR was 39.60 with a standard deviation of 7.74. The South Asian population has almost the same number of samples as the European population and both the distributions are observed to be more similar than any other pairs. The East Asian population has a very different distribution than all other populations. We found that almost 40.00% of the samples have CN in the range of 46 to 50 and the mean CN estimate and standard deviation are 44.60 and 6.77 respectively.

The DRAGEN LPA caller was able to phase the CN estimates for 47.06% (1507 of 3202) of the samples of the 1KGP dataset. Phased KIV-2 CN can be obtained in the case when we can determine the haplotype of the common SNPs utilized for phasing (see methods). We studied the phased CN estimates and their differences across all the populations. The distribution of phased CN differences are shown in **Figure 2B** and allele specific CN estimates are shown in **Figure 2C and 2D**. For EUR samples, the DRAGEN LPA caller was able to estimate phased CNs for 47.87% (303 out of 633) samples and the majority of all populations among all subgroups have the haplotype difference i.e. difference between two phased CN in the range from 0 to 10 (shown in **Supplementary Figure S7**). We also performed a two-sample Kolmogorov-Smirnov (KS) test by taking the distribution for EUR samples as reference to compare the distribution with other non-European groups. The high p-values with SAS and AFR (0.419 and 0.443 respectively) confirms the more similar distribution as shown in **Figure 2A** (also ECDF plot in **Supplementary Figure S3**). The distribution between EUR and AMR is observed to be significantly different (KS test p-value = 0.0365) this might be impacted by the lower number (186) of AMR (vs 303 EUR samples) samples have phased CN estimates. We also analyzed the haplotype difference among all seven subgroups of AFR population and the distributions are shown in **Supplementary Figure S8**.

### Correlation with previously known SNVs across *LPA*

Previous studies have utilized flanking SNVs to identify high risk KIV-2 CN numbers ^10,17,31^. We genotyped 24 such SNVs that have been previously reported ^10,24^. **Supplementary Table 6** contains the GRCh38 coordinates and the rsIDs of these SNVs. We performed Kolmogorov– Smirnov tests between copy number distributions in samples with the major allele in a homozygous state vs. samples with one or two copies of the alternate allele. **Figure 2E-L** shows detailed plots comparing CN numbers per population to observed SNV for a subset of the 24 SNVs investigated. **Supplementary Figure S9** provides the overview of remaining SNVs. This analysis confirms that KIV-2 copy number is correlated with several SNV alleles (**Supplementary Table 7**). However, this analysis also highlighted the disadvantages of using these SNVs predictively. At many of these sites, for example as shown in **Figure 2E & F** and **Figure 2K & L**, the copy numbers present in samples even with the homozygous alternate SNV allele, where the least variation of CN is expected, have CN ranges of greater than 35, while in the tightest groups (**Figure 2G and 2I)** the ranges are still ∼8 copies. Furthermore, we observed that most of the SNV investigated are often quite common (∼ 2.85%) thus not informative for the prediction of the CN KIV-2 itself (**Supplementary Table 7**).

Some minor alleles were also only present in a small number of samples, such as in **Figure 2I & 2J**, where 136 and 146 samples (∼4.20% and ∼4.60% of all samples) respectively contained at least one copy of the minor G alleles at rs10455872 (chr6:160589086) and rs41270998 (chr6:160599494). Others were absent from some populations, such as in **Figure 2G**, which shows that the minor allele T at GRCh38 coordinate rs41272114 (chr6:160585045) occurred in each population group except the East Asian group (EAS). The EAS population also does not contain the minor allele at the sites rs41272110 (chr6:160585140) and rs10455872 (chr6:160589086) as shown in **Figure 2H & 2J3**, while the minor alleles at the sites rs41270998 (chr6:160599494) and rs7760010 (chr6:160685571), as shown in **Figure 2I & 2L** respectively, are unrepresented in either the EAS or the SAS (South Asian) population groups. These results demonstrate a lack of portability of these alleles as predictors across populations.

These challenges, particularly the wide ranges of copy numbers present in samples homozygous for the minor predictive allele, illustrate the difficulty of SNV-based prediction of KIV-2 copy number to the degree necessary for confident detection of *LPA* alleles with increased risk of CVD, and shows the importance of more direct measurement of KIV-2 allele copy number. This highlights the need to assess the CN of KIV-2 directly using the DRAGEN LPA caller.

### Association with Lp(a) measurements across populations

Given the accuracy of the KIV-2 CN caller and the interesting patterns observed in the 1KGP data we next focused on different American populations. Here, we used a ∼30x WGS dataset of 3,006 randomly selected participants with self-reported populations: 1,004 European-American, 1,001 African-American from Atherosclerosis Risk in Communities (ARIC) cohort and 1,001 Hispanic-American from Hispanic Community Health Study / Study of Latinos (HCHS/SOL) cohort (see methods). Out of the 3,006, 2,375 had recent Lp(a) measurements or covariates.

Across the different populations we raise the question if the Lp(a) level is impacted from the SNV as reported for Europeans and as some SNV show impact across the 1KGP, or if there are other drivers at play across Hispanics- or African-American. To address this, **Figure 3A** shows the distribution of *LPA* CN among different populations. For these samples, the DRAGEN LPA caller was able to provide KIV-2 calls for all the samples and was able to report phased KIV-2 for 46.14% of samples.

**Figure 3:**
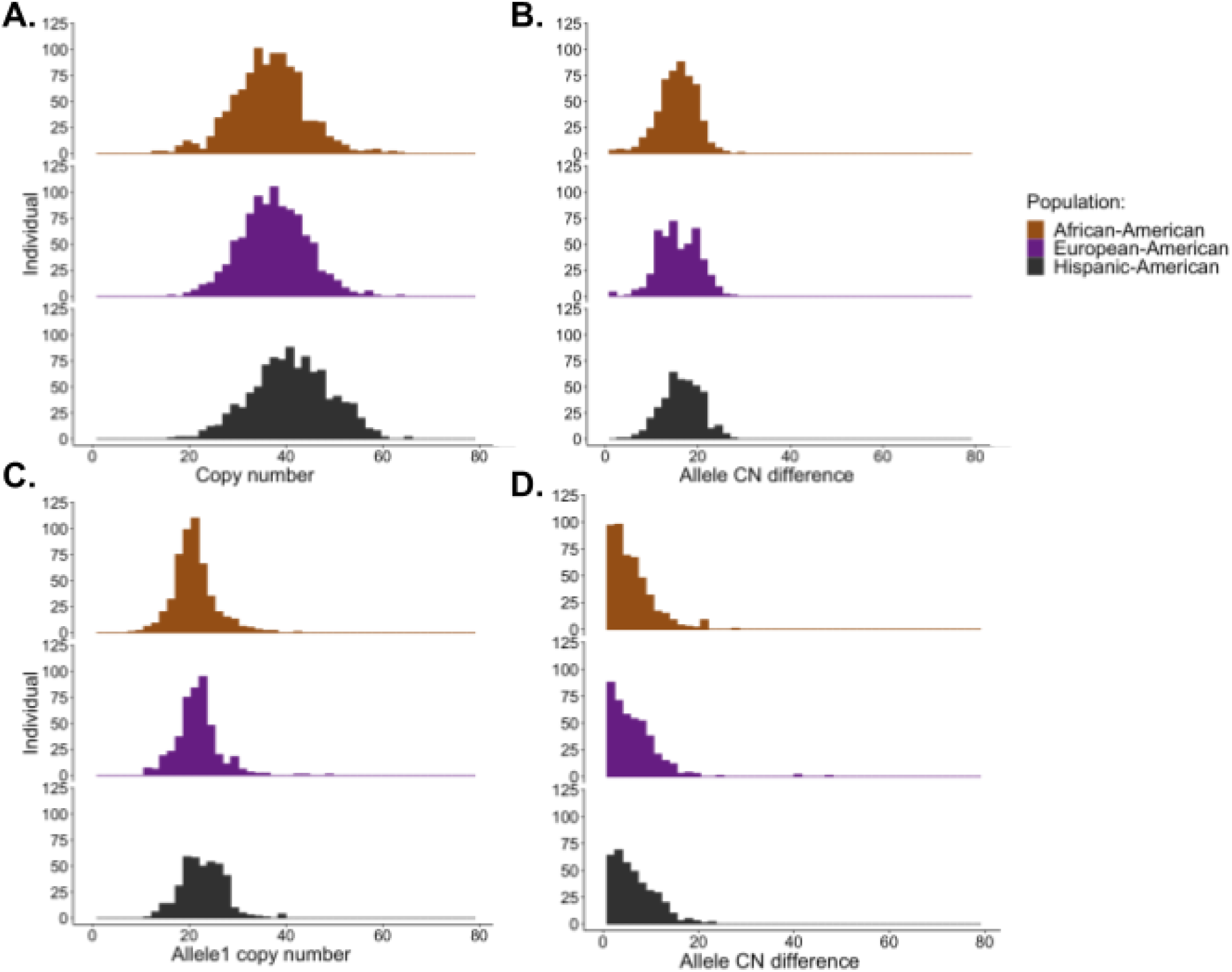
Overview of ARIC and SOL cohorts across the USA: **A)** Total KIV-2 copy number distributions in African-American, European-American and Hispanic-American populations. **B)** Haplotype difference between allele1 CN and allele2 CN among various populations. **C)** and **D)** distribution of Allele1 and Allele2 copy numbers. In each diploid genome with haplotype resolved KIV-2 calls, Allele1 is the longer allele and Allele2 is the shorter.

For European- and African-American samples, we observed a similar distribution with mean copy numbers of 37.50 and 36.50 and standard deviations of 6.75 and 7.50, respectively. Interestingly, we observed a different CN distribution among the Hispanic-American samples with a higher mean copy number (41.10) and higher standard deviation (8.01). We also observed that ∼50.00% of the Hispanic-American population has KIV-2 copy number in the range of 40 to 55 while ∼50.00% of the African- and European-American population are in the range of 34 to 42. To the best of our knowledge, there are no significant studies on CN analysis of the Hispanic-American population. Our results here show that the distribution patterns in the Hispanic-American population were observed to be different from the African- and European-American samples. The median CN estimate of Hispanic-American population was 41.00 (37.30 for European-American and 36.70 for African-American population).

We compared the results of ARIC samples with the similar analysis of 1KGP samples. For European-American samples, the CN distribution was consistent with EUR samples of 1KGP (mean CN 37.50 and standard deviation 6.86). However, we observed slightly different results for African-American samples with higher average CN estimates (36.50 vs 35.70) and also higher standard deviation (7.30 vs 6.68). This may reflect the smaller African-descended 1KGP subpopulation, with only 113 African-American samples as compared to 1,001 samples in the ARIC cohort.

The DRAGEN LPA caller was able to estimate allele-specific CN for 46.14% of samples (1,386 of 3,006). For African- and European-American populations, we computed the difference between the allele1 and allele2 CN estimates and observed that ∼90% samples have this difference in the range of 0 to 10 and standard deviations of 5.49 and 6.00 respectively. However, the Hispanic-American population showed a different distribution with ∼80% of the samples having allele CN estimate difference in the range of 0 to 10 and standard deviation of 4.48 (**see Figure 3B**). The distribution of allele-specific CNs are shown in **Figure 3C and 3D**. The CN estimates including allele specific estimations of all samples are given in the **Supplementary Table 8** and the phasing rates are given in **Supplementary Table 9**. The ECDF plots shown in **Supplementary Figure S10** and **S11** show the different distribution patterns of the Hispanic population in comparison to the African- and European-American populations.

We also performed the analysis for the correlation of CNs with 24 previously known SNVs across *LPA*, as performed for the 1KGP data. We performed Kolmogorov–Smirnov tests between CN distribution in samples with each major allele (i.e., homozygous reference) and samples with minor alleles i.e., heterozygous or homozygous alternate (shown in **Figure 4**). Our analysis on the ARIC and SOL cohort samples were also in agreement with 1KGP analysis. For the SNV site rs759106280 (chr6:160640854), all CNs corresponded to the major allele except one African-American sample (**Figure 4D**). There were no African-American samples with homozygous alternates at rs1853021 (chr6:160664263) and rs41272110 (chr6:160585140) SNV sites (**Figures 4E** and **4F**). We also observed another interesting pattern at rs200561706 (ch6:160594009) and rs1623955 (chr6:160664246), where no European-American samples contained the heterozygous states (**Figures 4G** and **4H**). Again, these results suggest a lack of marker transportability across different populations.

**Figure 4:**
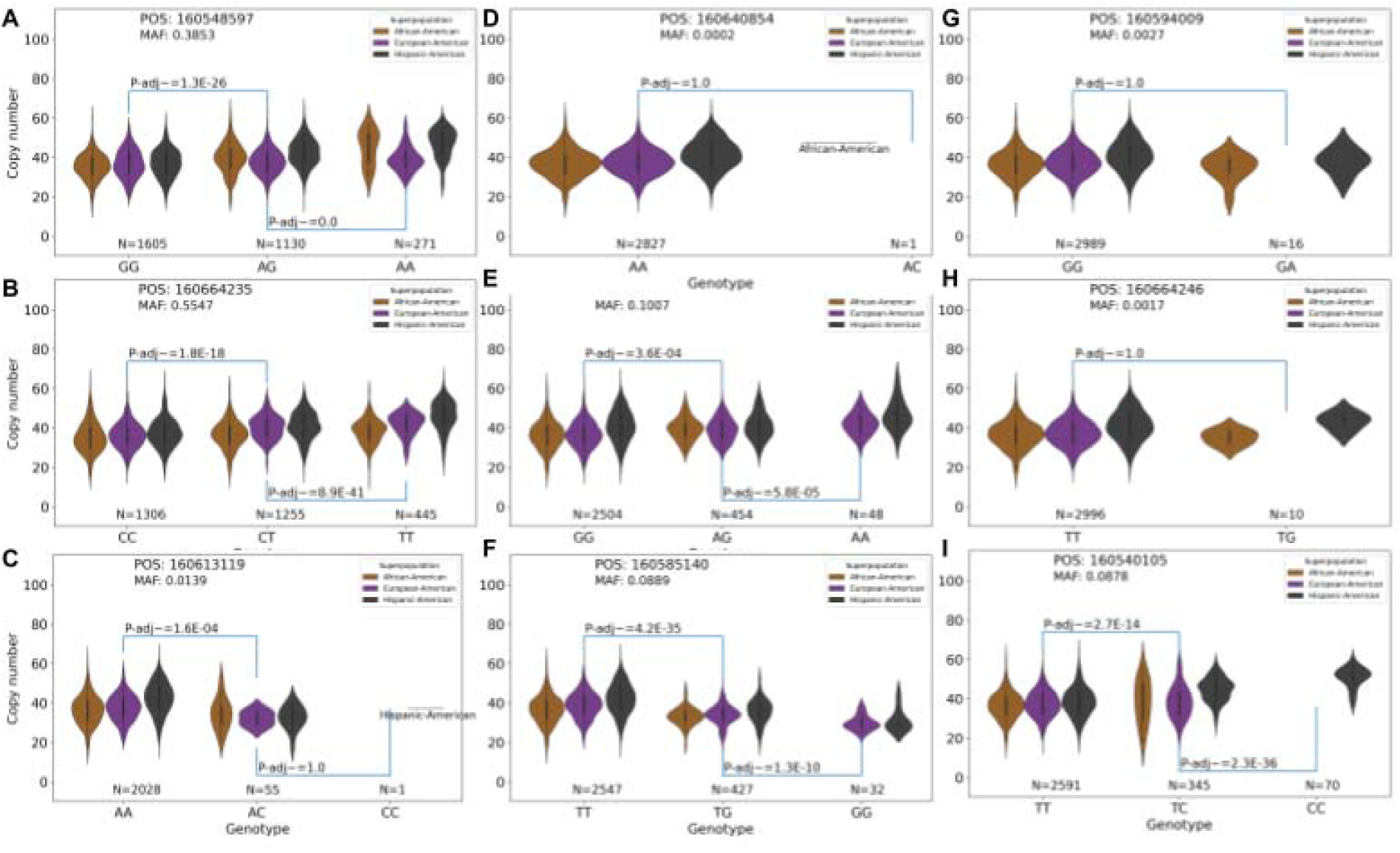
Comparison of SNV markers with CN states of KIV-2. Samples from ARIC and SOL cohorts were genotyped for 9 SNVs which have been reported as associated with different Lp(a) levels or copy number states. KIV-2 copy number was compared between groups of samples homozygous for the major (more common) or minor (less common) alleles of these SNVs, to test for association between SNV status and KIV-2 copy number. p-values were obtained by Kolmogorov-Smirnov tests and corrected for multiple testing by the Bonferroni method.

Next we compared the DRAGEN LPA caller estimation to protein measurements across all three American populations. We found that the Lp(a) levels in African-American (interquartile range (IQR) 70.10 (37.40-122.50)) were sparser as indicated by the wider IQR in comparison to European-American (IQR 23.10 (10.50-67.50)) and Hispanic-American (IQR 25.60 (9.80-66.20)) populations, which is consistent with published studies. Furthermore, we observed that CN quartiles were negatively associated with Lp(a) levels among African-, European- and Hispanic-Americans respectively (p-values: 4.03e-47, 2.15e-46, and 2.64e-42) and (partial R^2^ : 29%, 22%, and 32%). **Figure 5A** shows the relationship between predicted KIV-2 copy numbers by quartiles and the levels of Lp(a) across all 2,375 individuals for whom we had Lp(a) measurements, split by population. As expected, mean levels of Lp(a) were greater in the lowest quartile and attenuated in the higher quartiles. Of note, steeper declines of Lp(a) levels were observed in European- and Hispanic-Americans as compared to African-Americans. In all cases, we observed the previously reported correlation of high Lp(a) levels to low KIV-2 CN values using a linear model (see methods). This was highly significant for all three populations: African-American (p-value: 2.22e-45, partial R^2^ : 28%), European-American (p-value: 2.30e-56, partial R^2^ : 26%) and Hispanic-American (p-value: 1.07e-44, partial R^2^ : 34%). We also observed the inverse association between KIV-2 copy number and Lp(a) concentration for most KIV-2 length quartiles. These observations demonstrate the significance of allelic length, in addition to total KIV-2 copy number, in prediction of Lp(a) levels. **Figure 5B** highlights the effects of KIV-2 CN values with respect to clinical Lp(a) lipid thresholds of low (<75 nmol/L), medium (between 75 nmol/L and 125 nmol/L) and high (>125 nmol/L). Again, we can clearly see a difference of CN KIV-2 compared to the individual thresholds across all individuals highlighting the benefits of measuring the CN directly.

**Figure 5.**
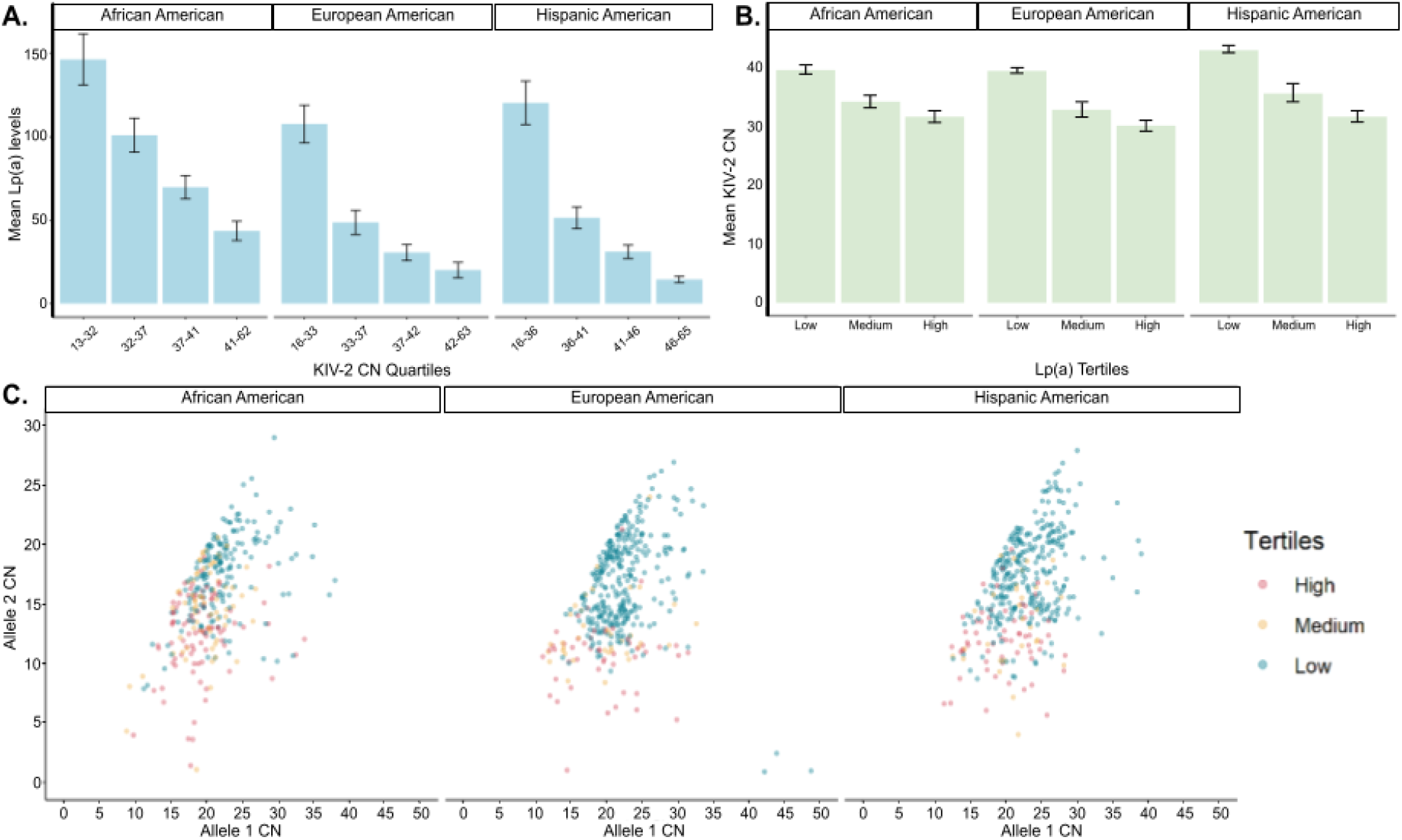
Association between KIV-2 CN estimates and Lp(a) measurement (in nmol/L): **A)** The mean Lp(a) level vs CN quartiles for African-American (p-value = 4.03E-47) and European-American (p-value = 2.15E-46) samples from ARIC cohort, and Hispanic-American samples from HCHS/SOL cohort (p-value = 2.64E-42). **B)** The mean LPA KIV-2 CN vs Lp(a) levels of three groups based on their diagnostic cutoff values of Lipid level i.e. <75 nmol/L as low, >125 nmol/L as high and in between 75 nmol/L and 125 nmol/L as medium. **C)** Scatterplot of allele length comparisons for KIV-2 CN and their impact on diagnostic cutoff values of Lipid levels per population. Allele 2 (Y-axis) is always reported as the smaller allele.

Next, we focused on the samples where we could obtain phased KIV-2 CN values to investigate the potential of phasing information. Here we compared phased allelic CN for allele 1 and 2, and identified similar relationships as observed with total KIV-2 copy number (**Supplementary Figure S12**). **Figure 5C** shows the relationship of the haplotypes per individual to corresponding Lp(a) level. Here the second haplotype is always reported as the smaller KIV-2 CN value of the two to enhance comparison across samples. It is interesting to note that we see slightly different patterns across the different populations. We can see the impact of different KIV-2 CN haplotype values per individual and the consequences for clinical cutoffs on lipid value reporting. This indicates that an extreme haplotype CN outweighs the other haplotype CN *LPA* value for lipid measurements, e.g., lower right in European-Americans (**Figure 5C**). Other outliers (i.e. high difference between haplotypes) do not show such a striking effect on the clinical lipid cutoffs potentially due to factors such as the previously reported decrease in *LPA* production from extremely short alleles^24^. We performed ROC analysis to measure the performance of allele 1 copy number (AUC 0.697), allele 2 copy number (AUC 0.839), and total copy number (AUC 0.846) for prediction of samples as likely having high Lp(a) (>125 nmol/L), suggesting that a model incorporating total copy number and the shorter allele copy number may result in high predictive power (**Supplementary Figure S13**). These results clearly highlight the importance of phased KIV-2 CN values to predict and study abnormal Lp(a) measurements and thus CVD risk.

In each population and for either allele, the difference between mean Lp(a) concentration for the first and second quartiles of KIV-2 allele length was the largest difference between contiguous quartile pairs.

Finally, we investigated the additional value of combining our CN KIV-2 values with the 24 SNVs investigated throughout this manuscript. Here we assess the association between Lp(a) concentrations and KIV-2 CN conditioning on SNV state (**Supplementary Table 10**). The African-American, European-American, and Hispanic-American populations each had non-zero minor allele frequency for 18, 17, and 19 of these 24 SNVs, respectively. We found that three, nine, and seven of these SNVs had significant effect sizes in the African-American, European-American, and Hispanic-American populations, respectively (p-values adjusted by Bonferroni method, alpha=0.05). In each population the association of KIV-2 copy number with Lp(a) concentration remained significant (p < 1e-25), suggesting an independent effect of CNs on Lp(a). Past studies have shown high Lp(a) levels and *LPA* SNVs are associated with CVD risk. Our study demonstrated that CNs had an independent effect of *LPA* SNVs and can have a reversed effect on Lp(a) levels, which may possibly alleviate the risk of CVD.

This highlights that CN values are more relevant than the previously postulated SNVs to predict Lp(a) levels in patients across the American populations. In contrast to SNV methods, our CN method shows a better performance and, more importantly, provides key insights into *LPA* even across different populations around the world.

## Discussion

Here we present the DRAGEN LPA caller, the first short read methodology to identify KIV-2 copy number. This KIV-2 CN caller only requires standard WGS Illumina sequencing and was developed as a component of DRAGEN targeted variant calling. This enables routine assessment of the copy number of the KIV-2 repeat in *LPA* and empowers research into the role of KIV-2 as a risk factor for CVD in single individuals or large cohorts. The DRAGEN LPA caller enables the phasing of KIV-2 CN alleles across 47.06% of individuals of 1KGP cohort and thus provides highly refined analysis of the role of allelic sizes on CVD risk. We show the accuracy of the DRAGEN LPA caller by comparison to 166 Bionano optically mapped genomes, with close correlation between the results of the two approaches when comparing either total KIV-2 copy number or phased allelic copy number. The ability of DRAGEN LPA caller to scale (runs within minutes) across sequencing data sets and provide detailed insights outmatches also available wet lab methods that are often laborious and do not provide comprehensive insights into the entire genome.

Traditionally the prediction of KIV-2 risk alleles have leveraged flanking SNVs that are often outside of the KIV-2 repeat and are in LD with certain risk CN allele size ranges ^32–34^. In this study, we investigated 24 SNVs postulated to provide key insights into KIV-2 CN numbers or to otherwise impact Lp(a) levels. Previous studies have shown that some of these SNVs are population-specific ^10,35,36^. In our study, we provide deeper insight, as our method allows direct comparison of KIV-2 copy number to allelic state for each SNV (**Figure 3 and 5)**, including across different populations. Thus, we were able to show the relationship of CN numbers and SNV states across different populations, including Hispanic-Americans and African-Americans. It is interesting to note that for some SNVs we indeed found significant correlation with CN numbers. However, the high transportability of direct *LPA* CN measurement makes this approach more broadly appropriate. Association of CN vs. SNV across Lp(a) further demonstrated that the CNs are a more portable predictor for Lp(a) numbers than SNV states across three populations. This is significant as it highlights the importance of our approach and its correctness. Thus, the DRAGEN LPA caller enables scalable population-wide analysis for the KIV-2 region based on widely available and cost-efficient Illumina whole genome data. The association study with Lp(a) measurements revealed a high R^2^ across non-European (African-American individuals (our R^2^ =0.28, p-value=2.22e-45 vs. R^2^= 0.038; p = .03)) population compared to most recent studies from UK Biobank ^37^. For a European-only population, Trinder et al showed a higher R^2^ value for Lp(a) than our study, likely because of the well characterized SNV. This again indicates that the CN KIV-2 is a better estimation across the human population overall. Furthermore, our analysis of the postulated SNV for Lp(a) revealed that they follow a bimodal population frequency distribution across the 1KGP. Thus, some SNVs appear to be too rare or too common to be diagnostically useful (**Supplementary Table 7**).

The accurate estimation of CN across KIV-2 is important to obtain estimations on the protein level of Lp(a) and with that on the associated cardiovascular risk levels per individual. Multiple questions about the genetic risk across populations have been postulated but not answered so far ^38^. With the DRAGEN LPA caller, we now have the methodology to reveal the CN across every Illumina-based WGS set. This method has already revealed interesting patterns around so-far understudied non-European groups (e.g. Hispanic-Americans). Overall, this study investigates KIV-2 repeat across 6,208 individuals from different populations. The DRAGEN LPA caller further demonstrates its ability to phase the CN into individual haplotypes, which enables a more detailed prediction of the expression values and isoform estimation of *LPA* directly. We demonstrated that this can be achieved in between 40-50% of the samples on average, with a slightly higher phasing rate in African-Americans (52.15%) than in European-Americans (45.72%) or Hispanic-Americans (40.46%). We further observed nearly bimodal KIV-2 copy number distributions in Hispanic-Americans, and to a lesser degree in European-Americans, although not in African-Americans. This is important given the insight that the two haplotype KIV-2 CN values are often different by 10 to 20 CN. **Supplementary Figure S14** also indicates a clear shift between the haplotypes, which may lead to a better explanation of Lp(a) overall values. Thus, the overall number of KIV-2 CN might not be the most reliable in assessing the CVD risk itself.

Implications of phased KIV-2 alleles remain to be determined in future studies. We speculate that phasing information will improve the cardiovascular risk prediction for many individuals that otherwise might not have correctly been diagnosed. This may be especially informative in cases where two haplotypes of KIV-2 have largely different copy numbers. We observed that the AMR population in 1KGP has the highest proportion (19.79%, 37 of 187) of samples with haplotype difference >10, 5.00% more than all other populations. For ARIC+SOL cohorts, Hispanic-Americans were shown to have a higher proportion (19.75% vs 13.41% and 15.03% for African- and European-American respectively) of samples with haplotype difference >10. Here the genome-wide or averaged CN could hinder the detection of a risk allele (i.e. low CN of KIV-2).

We observed a significant correlation between the phased allelic copy number and Lp(a) measurements, but it remains to be seen based on e.g. RNA experiments to what extent the phased CN values improve prediction, given complicating factors such as splicing variation.

Overall, we demonstrate that the DRAGEN LPA caller provides highly accurate KIV-2 CN calling even with phasing information for individuals. This enables the assessment of KIV-2 and *LPA* across millions of data sets and represents an inexpensive way to assess an important factor in cardiovascular risk utilizing traditional Illumina sequencing.

## Methods

### CN confirmation with optical mapping technology

Orthogonal confirmation is a key aspect of this study to show the accuracy of both total and allelic copy number calling. Therefore, we curated a set of KIV-2 copy number calls from publicly available long-contig data, using optical maps from Bionano Genomics.

Bionano optical mapping is a powerful tool to resolve this type of structural variation, as it is highly sensitive to large structural rearrangements ^28^. We used a set of 12 public Bionano genomes and generated an additional set of 154 Bionano genomes (see **Supplementary Table 1**) to measure the repeat lengths in this region. However, despite the reliability of optical maps, the KIV-2 locus proved challenging because it requires measurements for DNA that span the entire length of the repeat. Of the 332 alleles measured by optical mapping, 15 did not span the region and could not be confidently reported. Additional genomes where neither allele was spanned were dropped from analysis. This difficulty appears to stem directly from the unique structure of this VNTR, which may be hundreds of kilobases in length and does not contain the recognition sites used for Bionano mapping. However, the result of our Bionano variant curation is 317 high-confidence insertions/deletions that we converted to KIV-2 copy number.

### Estimating KIV-2 Copy Number

The DRAGEN LPA caller uses Illumina whole-genome sequencing (WGS) to measure the copy number for KIV-2 (see **Figure 3A and 5A**). The KIV-2 region includes six copies of an (∼5,500 bp region) in both the GRCh37 and GRCh38 human reference the DRAGEN LPA caller counts reads aligned anywhere within the KIV-2 region, which includes six copies in the GRCh37 or GRCh38 human reference genomes. Reads are then counted from 3,000 additional regions for normalization, each 2,000 bases in length. For quality control, the median absolute deviation (MAD) score is calculated and used to flag samples with high variation in coverage, with a maximum MAD threshold of 0.11. In these cases, KIV-2 copy number will still be reported but potentially with lower accuracy.

Read counts for all regions are normalized by region length, then by GC content using LOWESS regression ^28,39^. This smoothing method utilizes read counts from the set of normalization regions with their GC contents to predict the best adjustment for the read coverage of the KIV-2 region via a generalized linear model. The resulting normalized KIV-2 coverage metric, representing the number of copies of the entire KIV-2 region as represented in the reference genome with the KIV-2 copies it contains. The normalized KIV-2 coverage is then scaled by six to represent instead the number of copies of the KIV-2 repeat unit in the reference genome. This scaled value is the total copy number of KIV-2 in the sample.

### Estimating Allelic KIV-2 copy number

An important task for KIV-2 copy number calling is differentiation of allelic copy numbers, as total copy number does not directly indicate the lengths of the two alleles present in the diploid human genome. We are able to do this because, as described above, we identified two linked SNV sites at positions 296 and 1,264 within the repeat unit that, if present in a KIV-2 haplotype, occur in every copy of the repeat unit in the haplotype (T>G at chr6:160630428/160635977/160641520/160624884/160619338/160613786 and C>G at chr6:160620306/160625852/160631396/160636945/160642488/160614754, GRCh38). These two sites occur in linkage and are sufficiently common to appear heterozygously in 47.00% of the 1KGP cohort samples (**Figure 3C)**, with high frequencies of the useful heterozygous case in each population group.

To calculate allelic copy numbers, the DRAGEN LPA caller measures the proportions of reads containing the reference and alternate alleles at each of these two marker sites. Individual alleles can be predicted when this ratio is neither 0.00% nor 100.00%. If a minimum depth of ten reads supports both reference and alternate alleles, the resulting proportions are used to scale the total copy number measurements from all reads in the region into the copy number from the allele containing the marker SNV alternate alleles and the allele with the reference case at those marker sites.

The DRAGEN LPA caller is able to make a total KIV-2 copy number determination in 100.00% of samples with high-quality WGS and allelic copy number determination in ∼47.00% of the 1KGP samples.

### Association study

Lp(a) levels were inverse-normal transformed prior to the analysis by population groups. Linear regression was used for the ARIC cohort while survey linear regression was performed in the HCHS/SOL cohort. SOL participants were recruited by using a 2-stage probability sample design; survey linear regression accounts for the 2-stage stratified sampling and clustering of participants within sampling units. It is established and recommended by SOL ^40^. The model studied the effect of CN (total copy, allele 1, and allele 2) on Lp(a) levels adjusting for age, sex, and center. Linear trend test was performed for the quartile groups of *LPA* CN by allocating them into four ordinal groups, treating them as continuous. We additionally conducted a linear regression to analyze the association between *LPA* CN and Lp(a) levels, conditioning on 24 SNVs respectively. The statistical significant level is defined as p-value < 0.005, accounting for three types of CNs and populations analyzed.

The Atherosclerosis Risk in Communities (ARIC) is a prospective cohort study conducted on four communities: Forsyth County, North Carolina; Jackson, Mississippi; northwestern suburbs of Minneapolis; Minnesota, and Washington County, Maryland. The aims were to identify risk factors for subclinical atherosclerosis and cardiovascular disease (CVD). In 1987, it recruited 15,792 adults, ages ranging from 45 to 64. The study is still ongoing with visit 9 being the latest. Data collection was through phone interviews and in-person examinations ^41^. Lp(a) levels were measured at visit 4 through applying a double-antibody ELISA approach. The reliability was evaluated from diving between-person variance by total variance^41,42^. Values were standardized based on a conversion equation from a comparison between visit 4 samples measured in 2 assays in 100 samples from a whole Lp(a) distribution^43^.

The Hispanic Community Health Study / Study of Latinos (HCHS/SOL) is a prospective cohort that studied the prevalence of CVD in four communities: Bronx, New York; Chicago, Illinois; Miami, Florida; and San Diego, California ^40^. Between 2008 to 2011, the study randomly recruited 16,415 adults with age between 18 and 74 using a stratified two-stage probability method ^44^. Participants had on-site visits at baseline and received phone calls for follow-up. Lp(a) levels were measured at visit 1 via Monoclonal Antibody-based ELISA that can capture Lp(a) antigens using two monoclonal antibodies. This method can recognize all apo(a) isoforms on equimolar basis (nmol/L). Four control samples from single donors were used to assess precision. Analyses were performed in duplicates where samples are re-analyzed if the coefficient of variation surpasses 10.00%.

## Conclusions

Overall, we demonstrate that the DRAGEN KIV-2 caller provides highly accurate KIV-2 CNV calling even with phasing information for individuals. This enables the assessment of KIV-2 and LPA across millions of data sets and represents an inexpensive way to assess an important factor in cardiovascular risk utilizing traditional Illumina sequencing.

## Data availability

The 1000 genomes Whole genome is accessible from: http://ftp.1000genomes.ebi.ac.uk/vol1/ftp/

ARIC and SOL samples are accessible over dbGap and request from: https://sites.cscc.unc.edu/aric/distribution-agreements https://sites.cscc.unc.edu/hchs/

Software is available from Illumina at https://www.illumina.com/products/by-type/informatics-products/dragen-bio-it-platform.html

## Acknowledgements

The authors thank all the participants in SOL and ARIC studies.

## Authors’ Contributions

All authors contributed to manuscript writing and enhancing. XC, ME and JB developed the software. SB and JB performed the analysis for KIV-2. NN and BY performed the association study for ARIC and SOL. ME and FS designed the study.

