## Supplementary Figures for "Identification of allele-specific KIV-2 repeats and impact on Lp(a) measurements for cardiovascular disease risk"

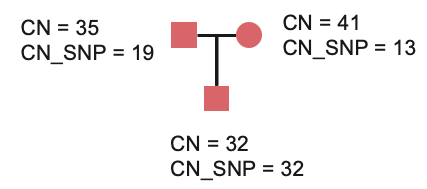

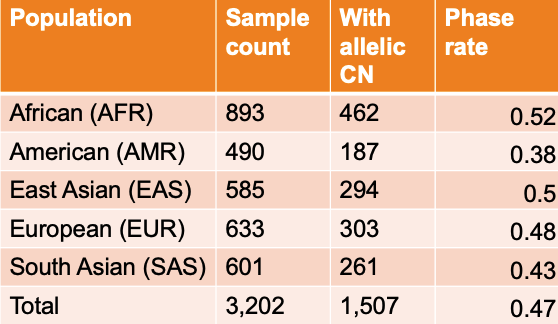

***Supplementary Figure S1 : Allele-differentiating SNV sites identified in 1KG samples****.* ***A.*** *Variant-predicted copy number compared against total inherited copy number. If the SNV allele in question occurs in every repeat unit from exactly one KIV-2 haplotype in each parent, the SNV read fraction in the parents must be reflected in the offspring KIV-2 copy number. Here, the father contains 35 total copies of KIV-2, with 19 of those copies containing the SNV of interest. The mother contains 41 total copies, with 13 containing the SNV. The SNV of interest is present homozygously in the offspring, with total copy number of 32 resulting from 19 paternal copies and 13 maternal copies.* ***B.*** *The frequency of heterozygous genotypes for the markers found for different populations. These sites can be used to differentiate allelic copy numbers wherever present heterozygously and are usable in 47% of the 3,202 samples from 1KGP.*

| A  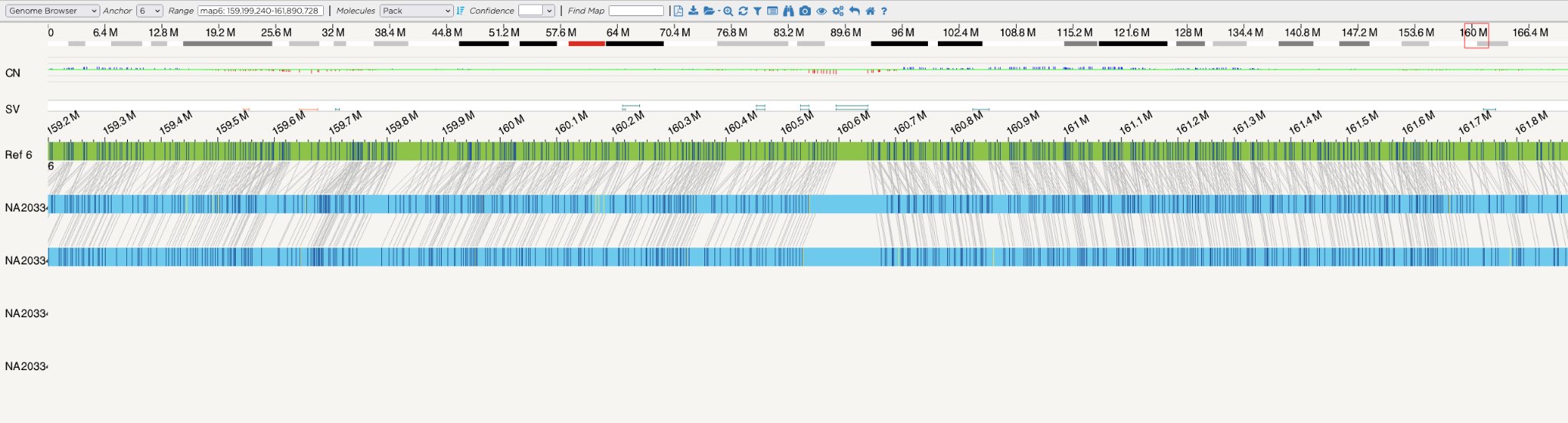 | |
| --- | --- |
| B  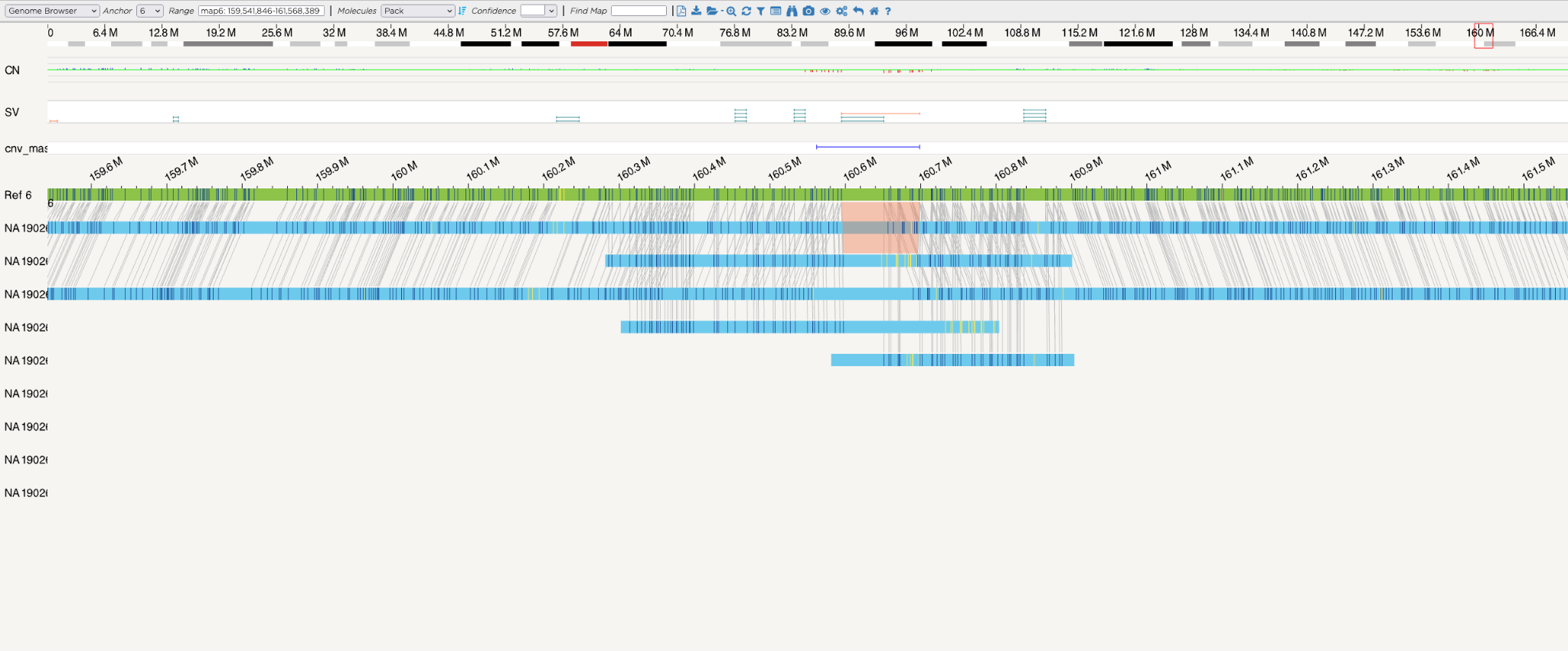 | |
| C  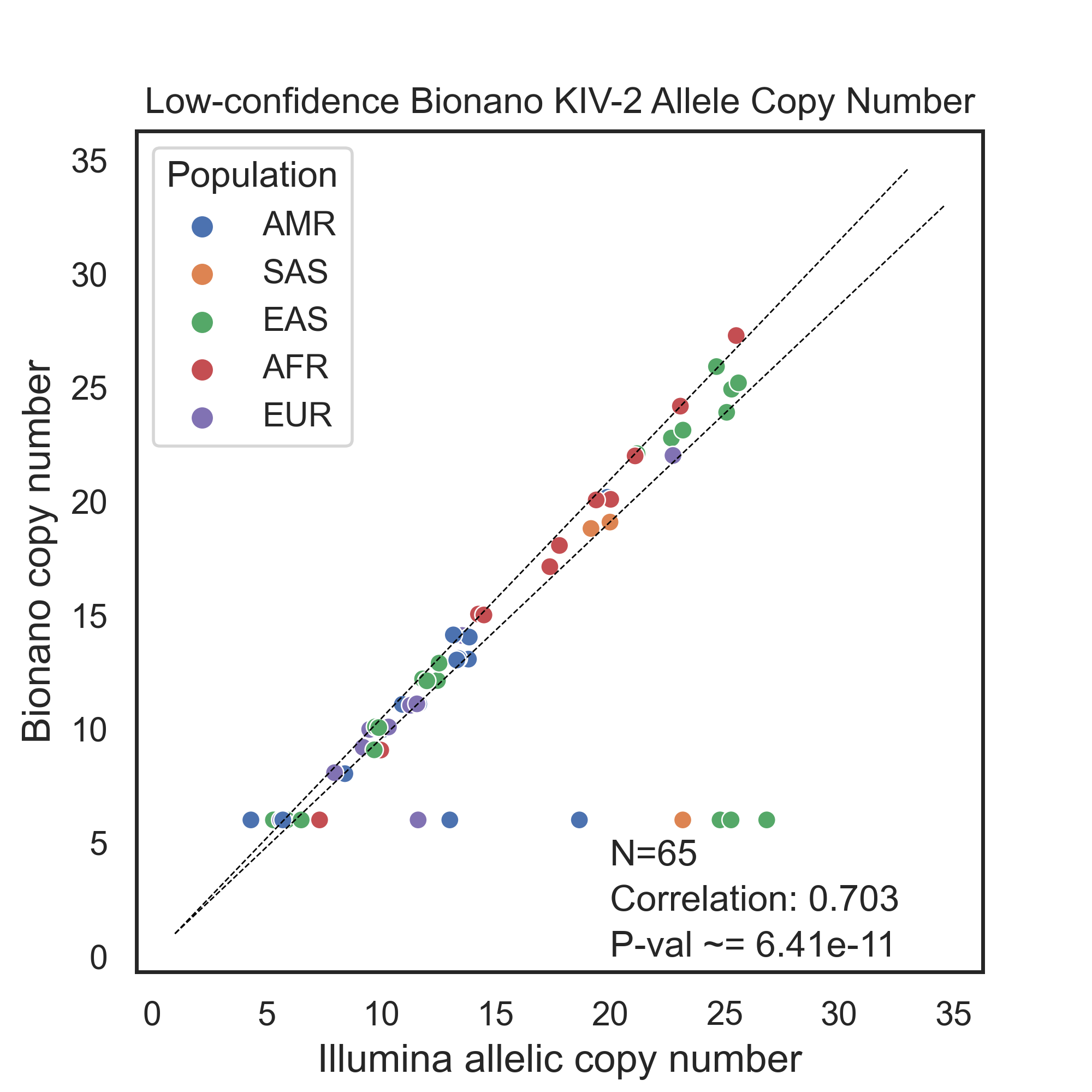 | D  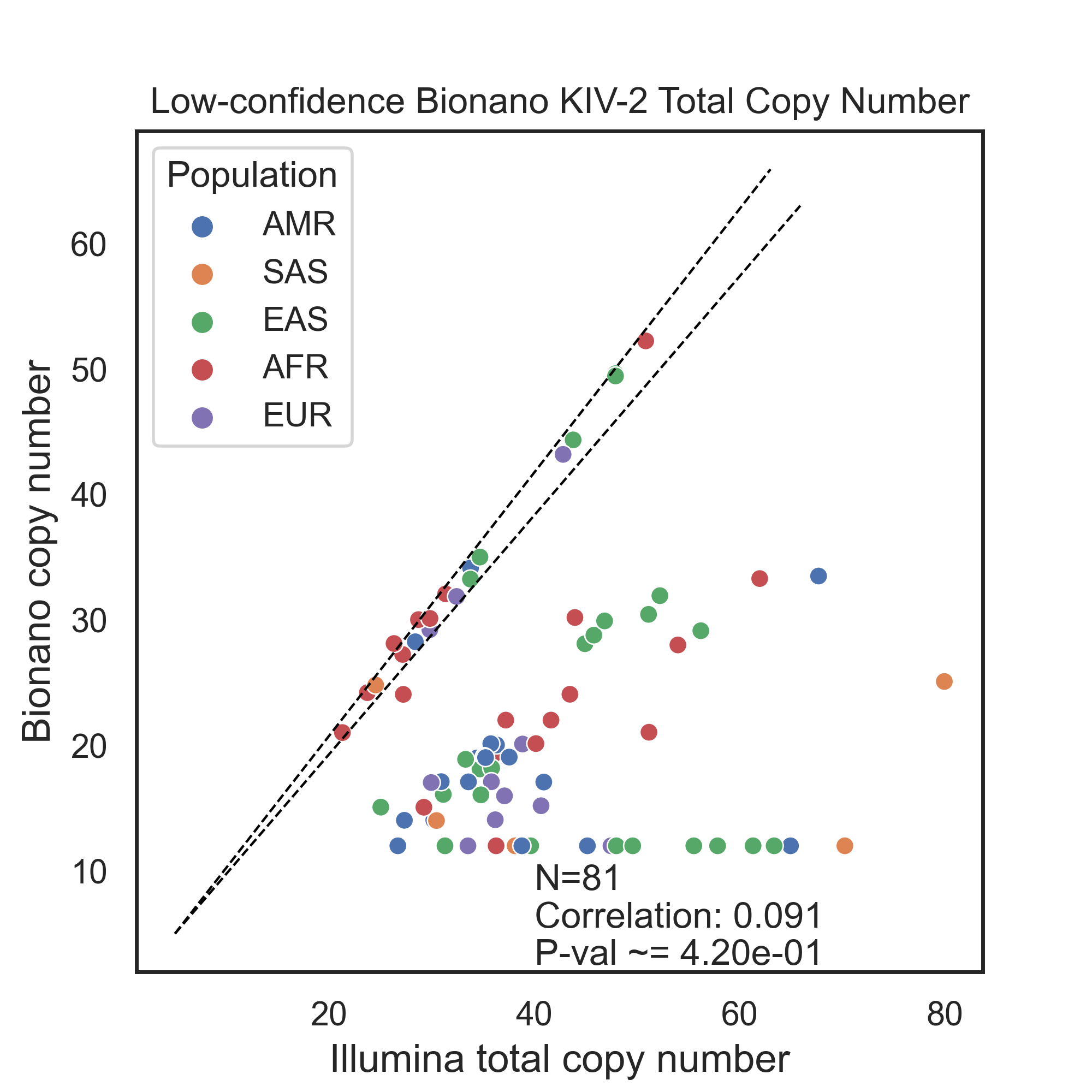 |
| ***Supplementary Figure S2. Bionano optical mapping challenges across* LPA*. A.*** *Successful assembly across the LPA locus generates two continuous haplotype assemblies or contigs, where multiple long DNA molecules are imaged and joined together to fully span the repeat KIV-2.* ***B****. In some cases, the length and high homology of the repeat leads to failed partial assemblies with non-standard ploidy, and potentially deleterious effects on structural variant calls from the failed assemblies.* ***C.*** *Comparisons between low-confidence Bionano-predicted KIV-2 allelic copy number and DRAGEN-predicted KIV-2 copy number, where assembly breaks and missing calls from Bionano assemblies suggest potential inaccuracy.* ***D.*** *Comparisons between low-confidence Bionano-predicted KIV-2 total copy number and DRAGEN-predicted KIV-2 copy number, where assembly breaks and missing calls from Bionano assemblies suggest potential inaccuracy.* | |

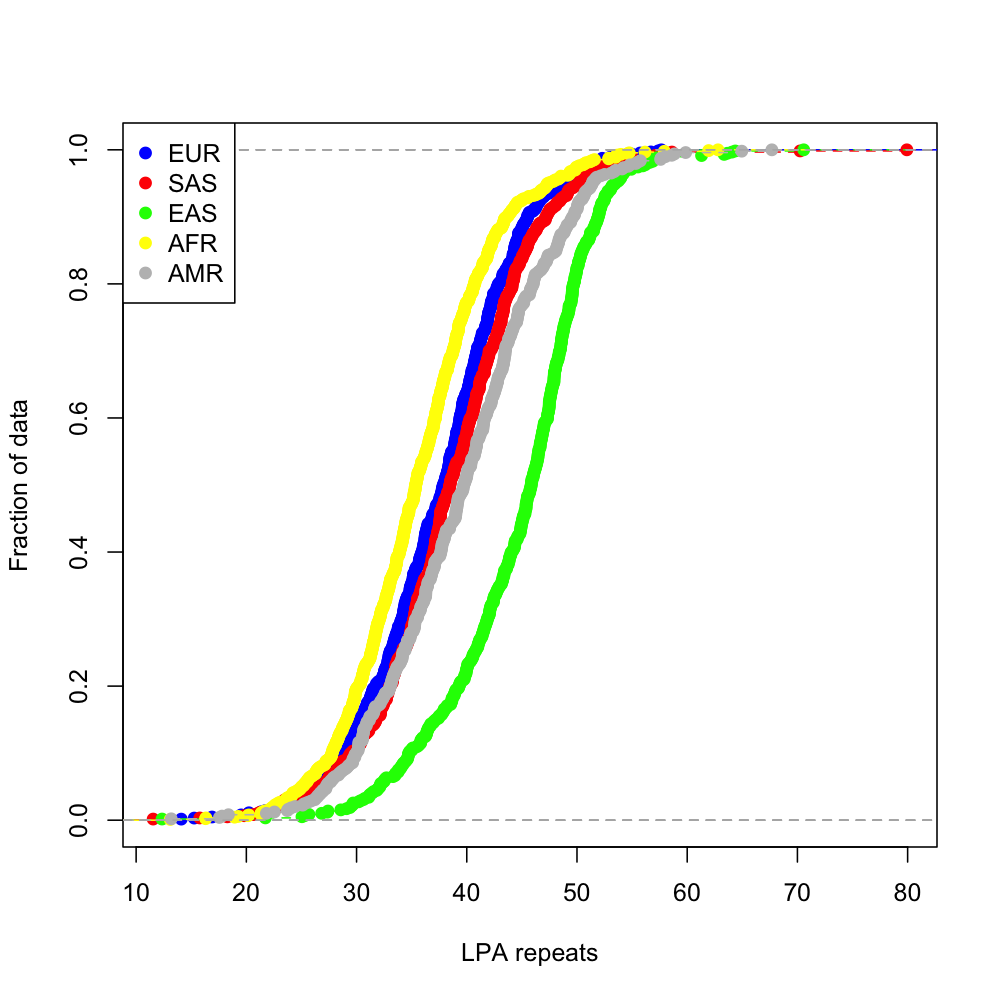

**Supplemental Figure S3**: Empirical Cumulative Distribution Function plot for CNV estimate distribution of all ethnicities in 1KGP dataset.

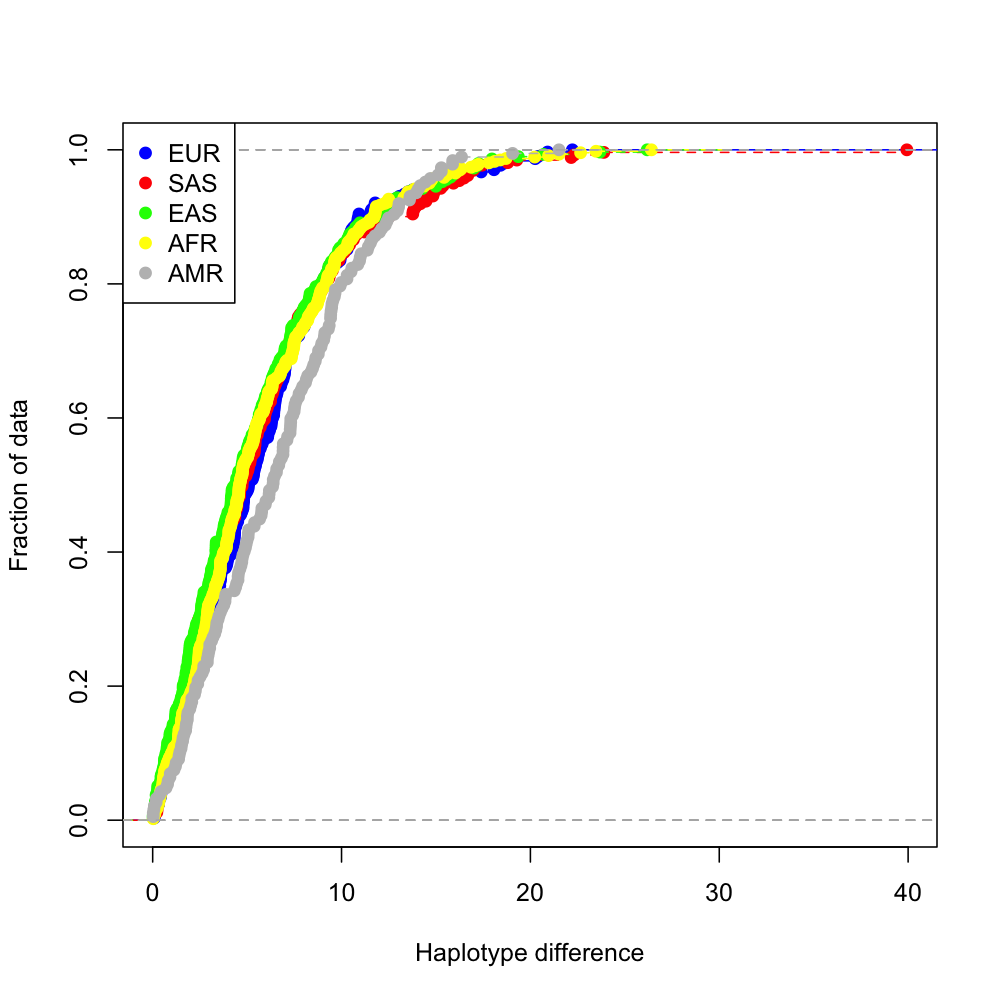

**Supplemental Figure S4**: Empirical Cumulative Distribution Function plot for phased CNV estimate difference i.e., abs (allele1_CN - allele2_CN) distribution of all ethnicities in 1KGP dataset.

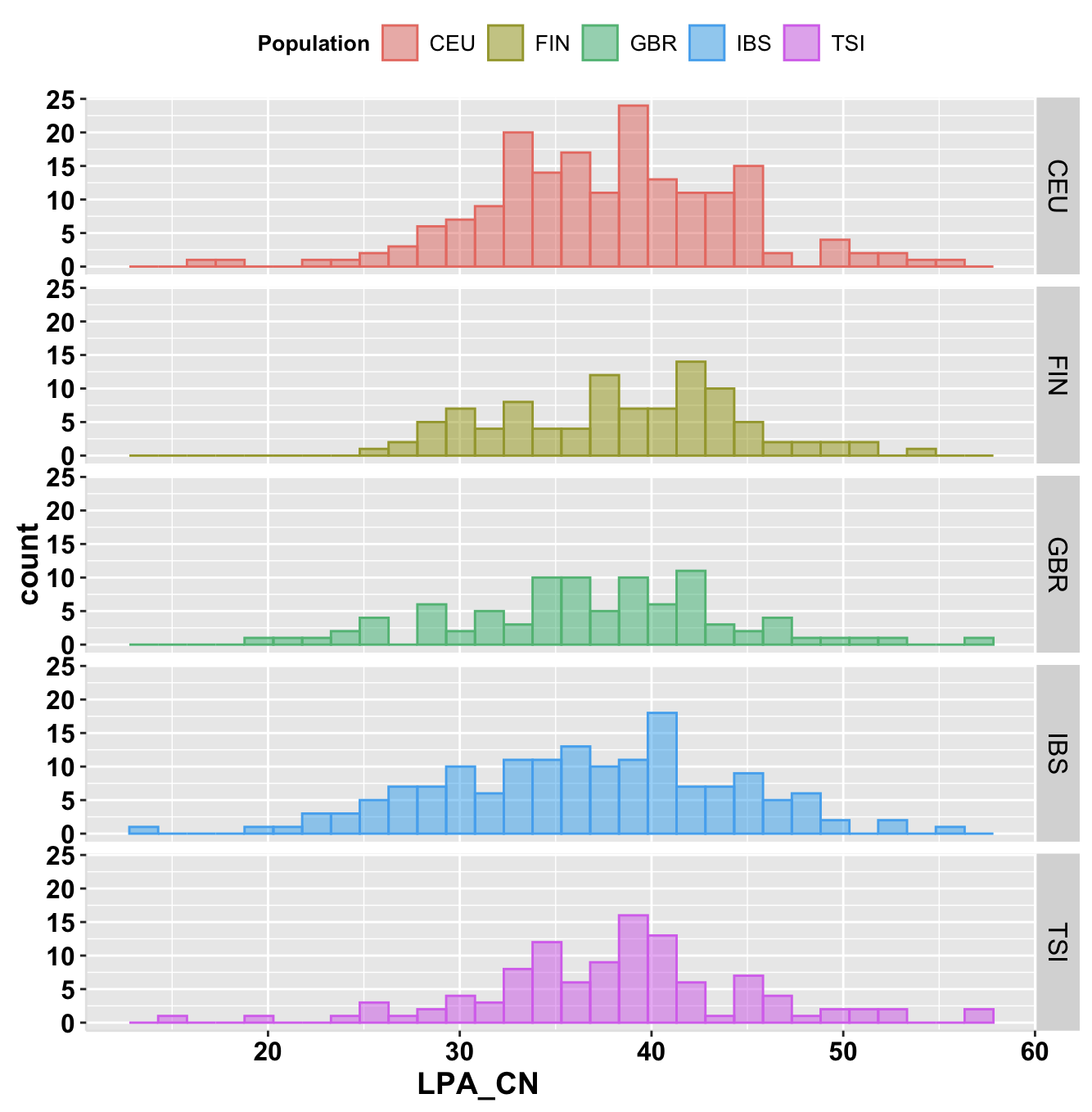

**Supplemental Figure S5**: KIV-2 CNV distribution among EUR subgroups (179 CEU, 99 FIN, 91 GBR, 157 IBS and 107 TSI)

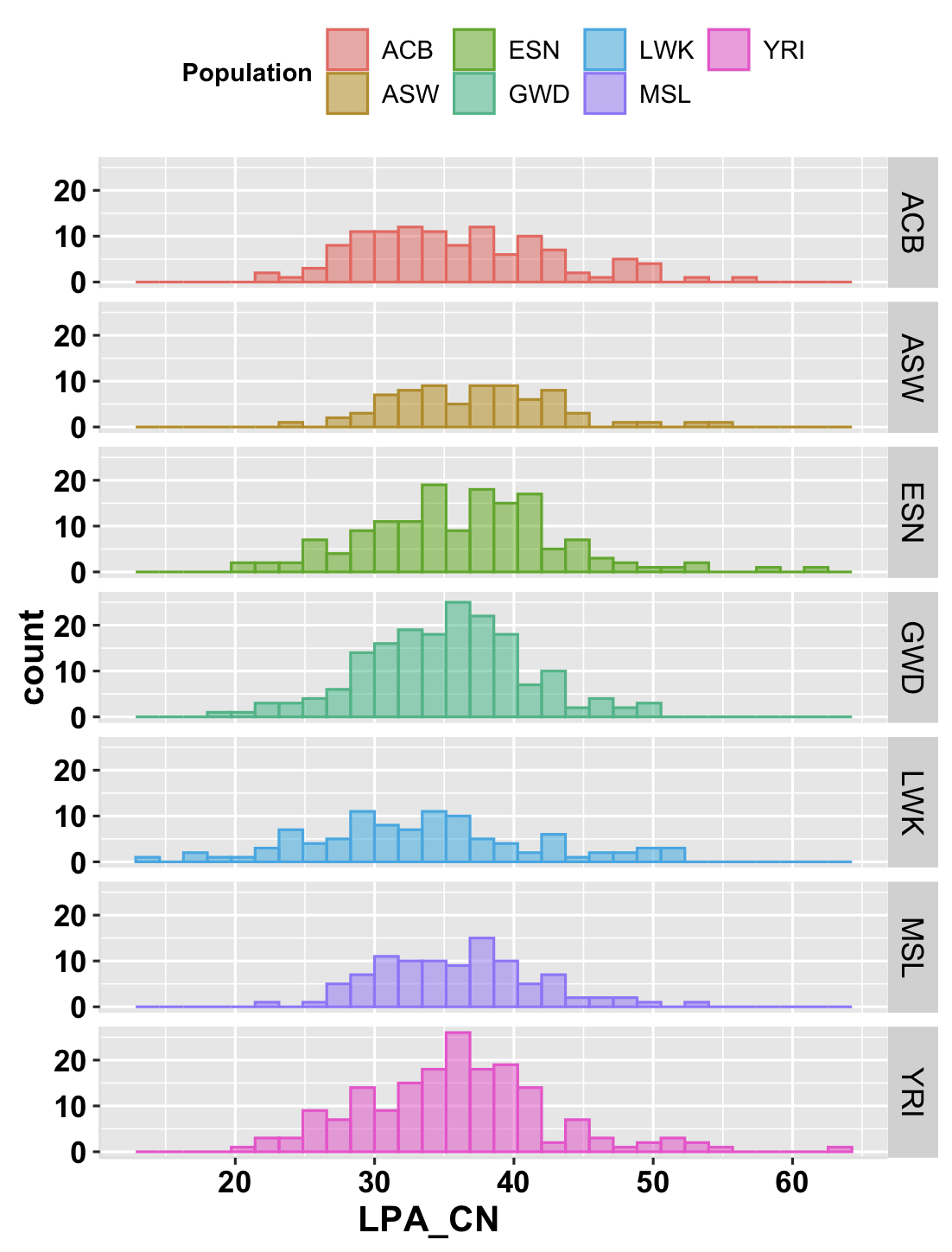

**Supplemental Figure S6**: KIV-2 CNV distribution among AFR subgroups (178 YRI, 99 MSL, 99 LWK, 178 GWD, 149 ESN, 74 ASW, 116 ACB) in 1KGP dataset.

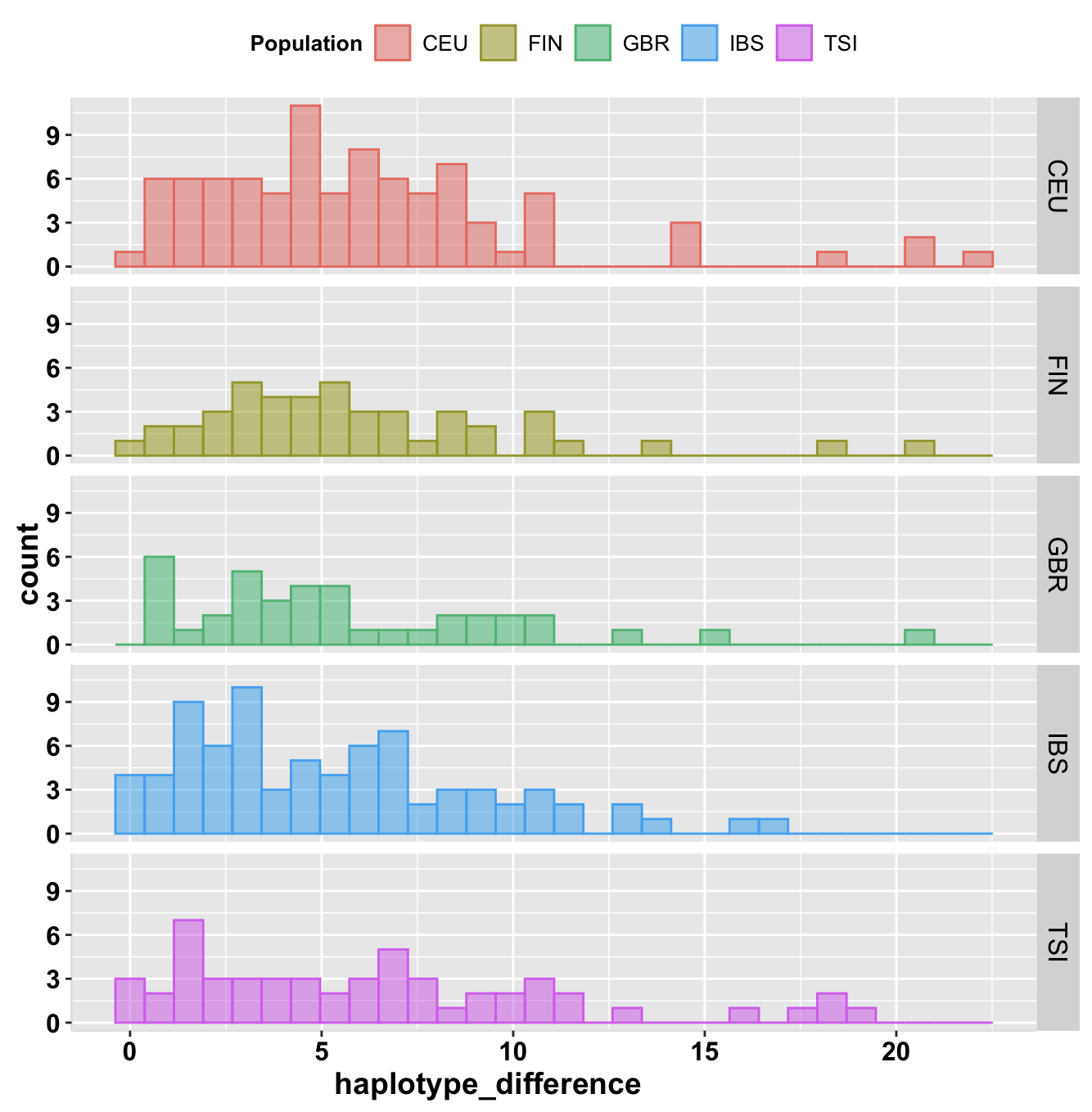

**Supplemental Figure S7**: KIV-2 CNV haplotype difference distribution among EUR subgroups (total 303 sample) in 1KGP dataset.

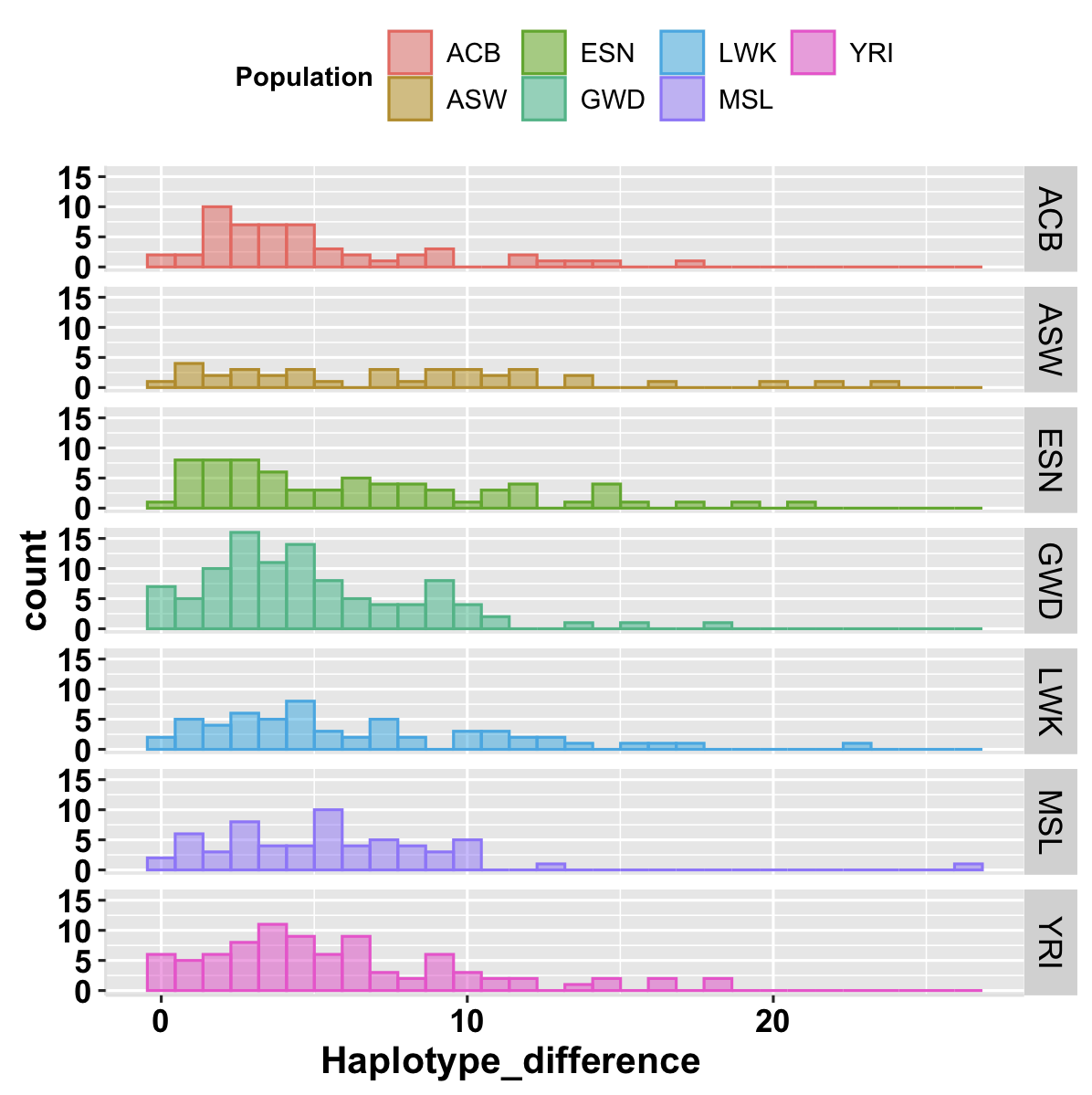

**Supplemental Figure S8**: KIV-2 CNV haplotype difference distribution among AFR subgroups in 1KGP dataset (total 462 sample).

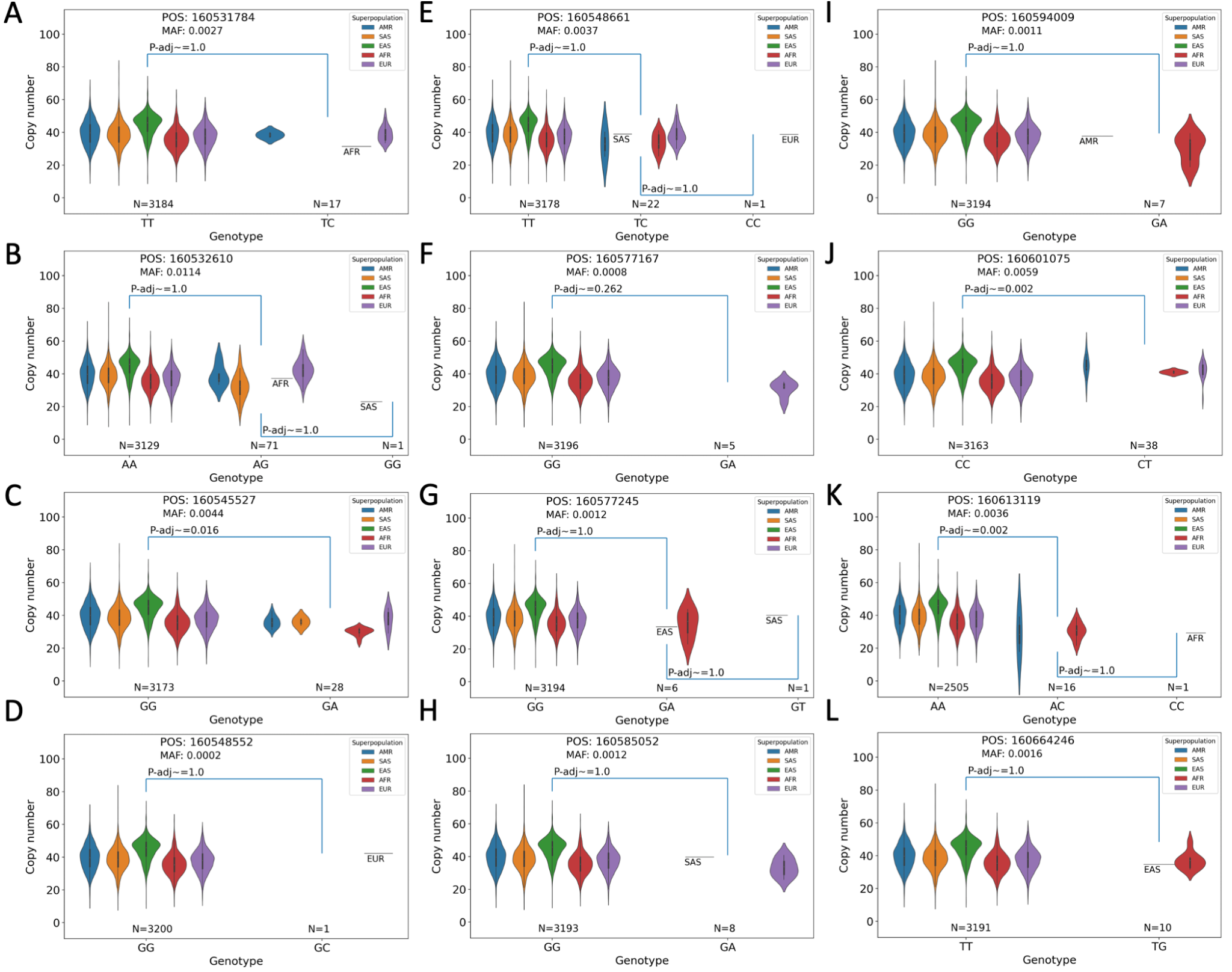

**Supplemental Figure S9**: Comparison of SNV markers with CNV states of KIV-2 in 1KGP.

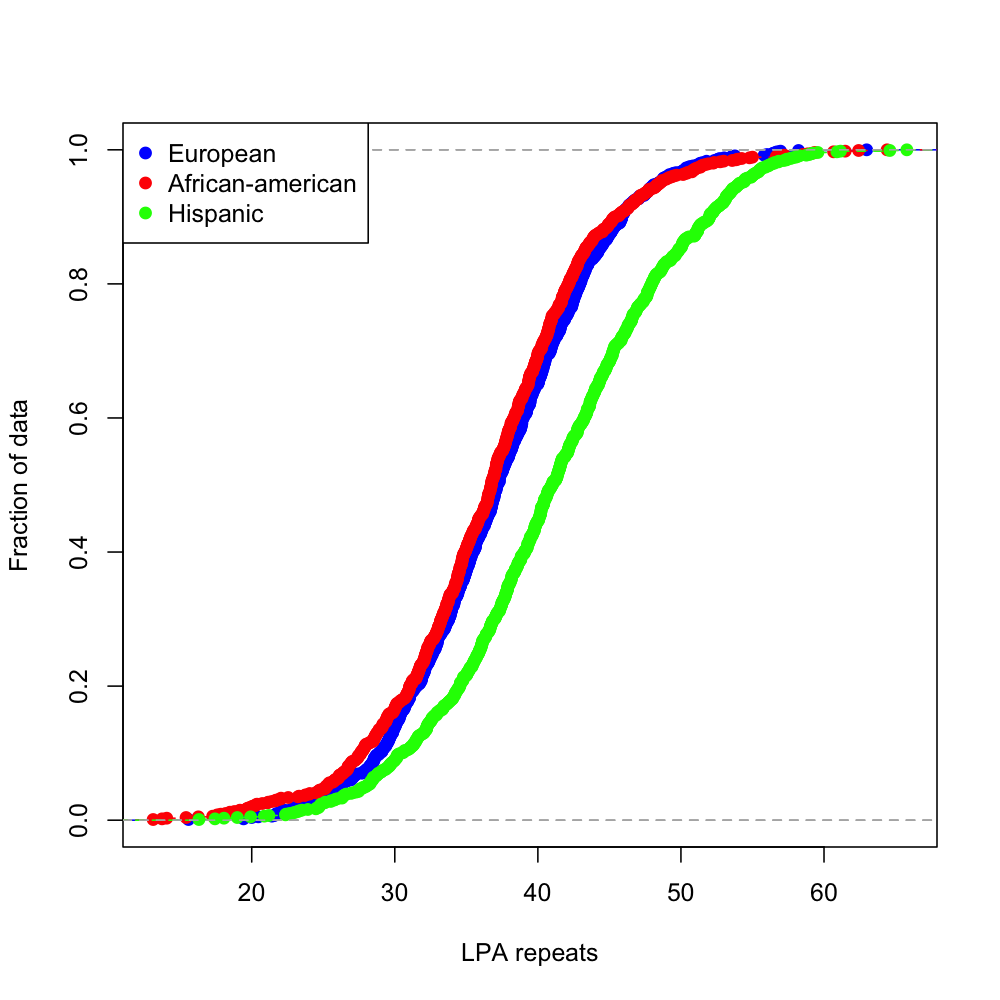

**Supplemental Figure S10**: Empirical Cumulative Distribution Function plot for CNV estimate distribution of all ethnicities in ARIC and HCHS/SOL cohort dataset.

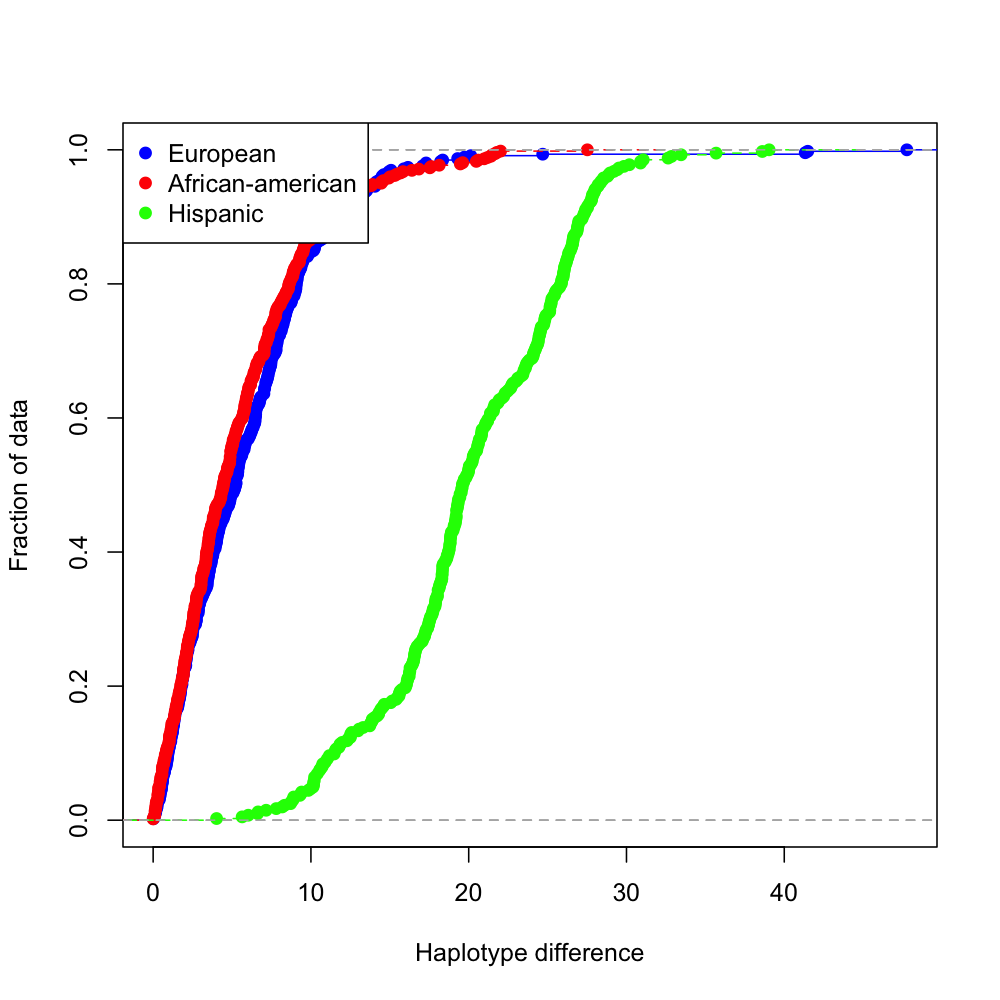

**Supplemental Figure S11**: Empirical Cumulative Distribution Function plot for phased CNV estimate difference i.e., abs (allele1_CN - allele2_CN) distribution of all ethnicities in ARIC and HCHS/SOL cohort dataset.

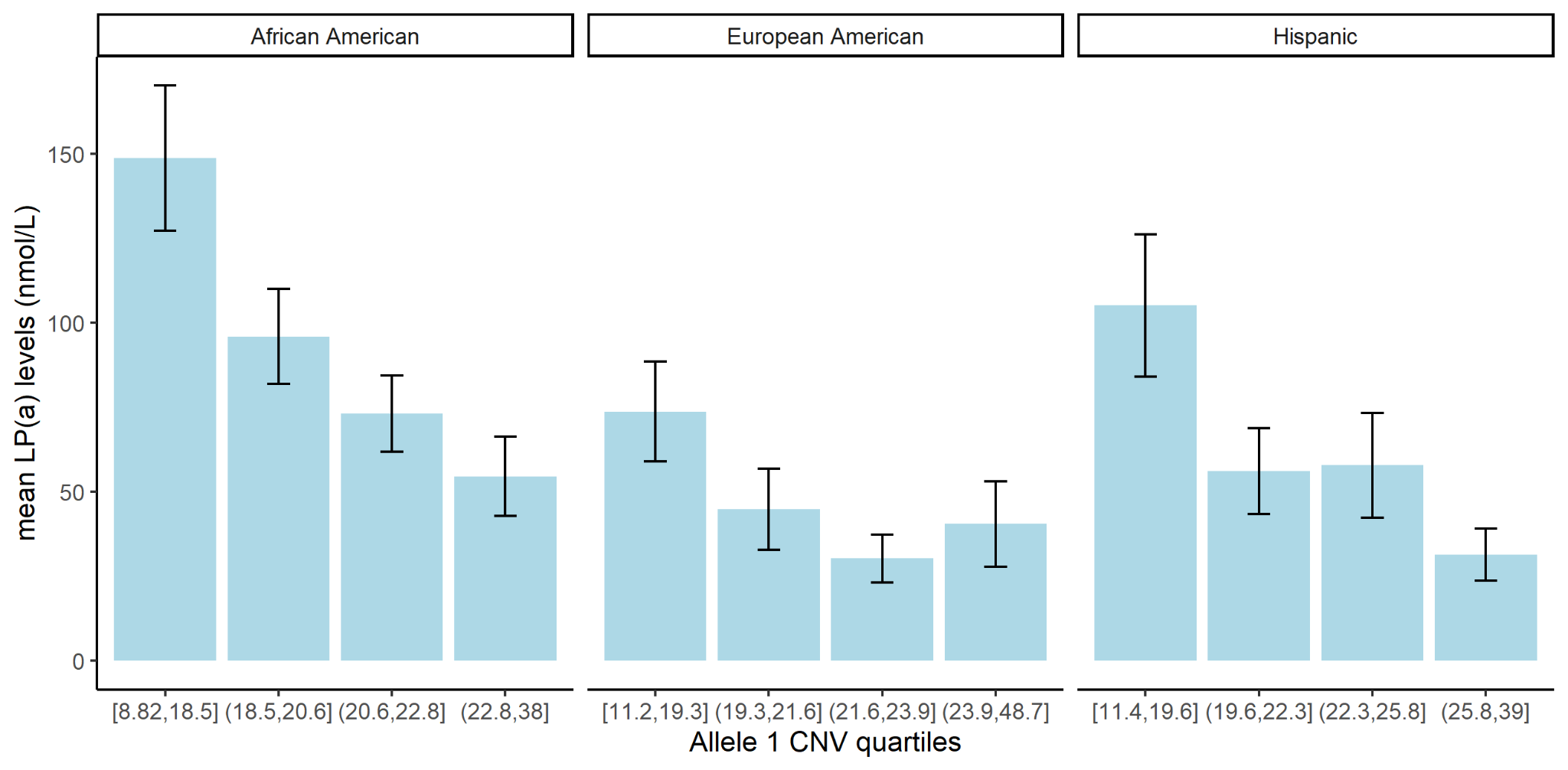

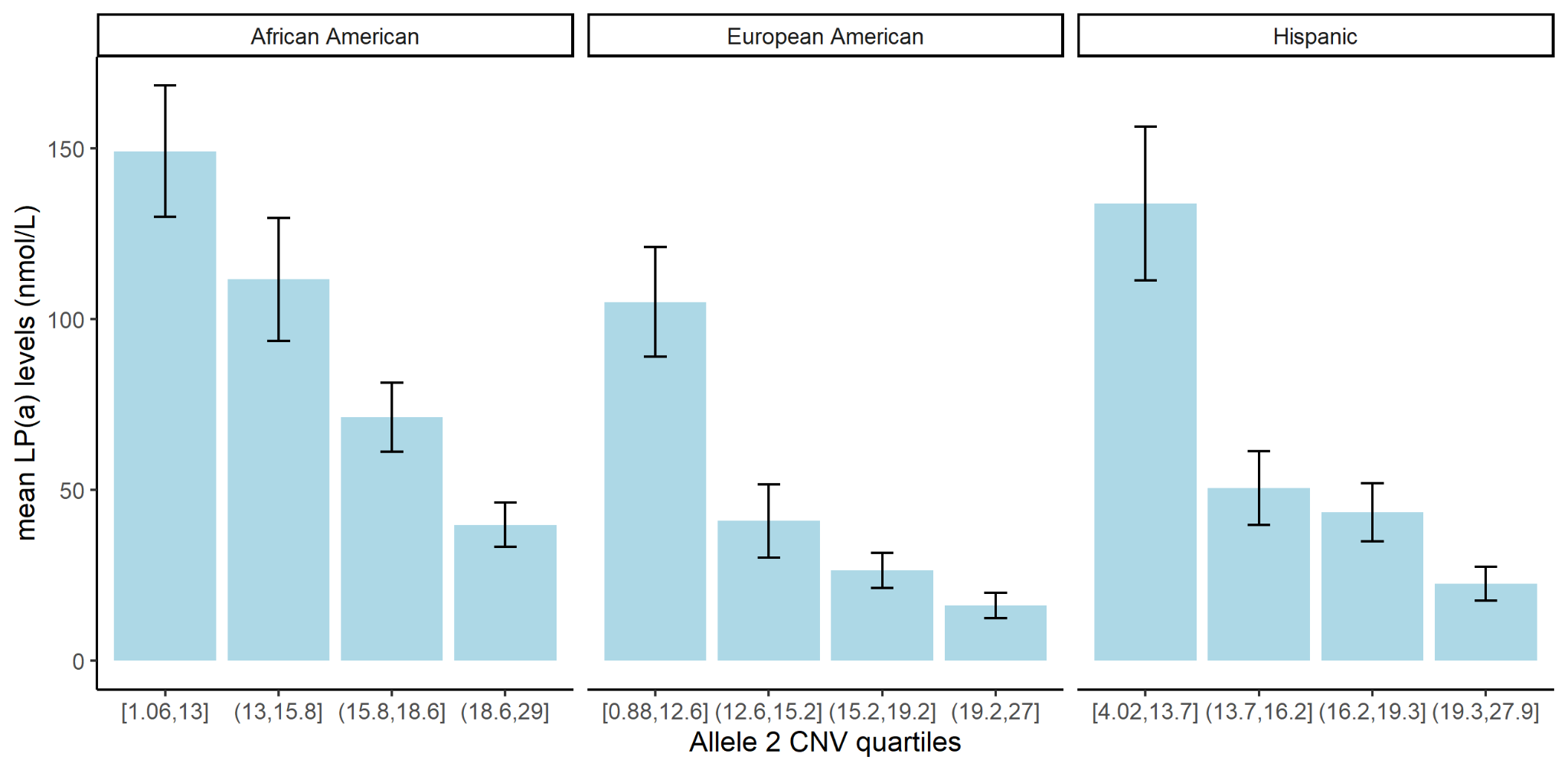

**Supplementary Figure S12: Allele length quartile associations with Lp(a) concentrations.**

**A**. Allele 1 (longer allele) CN quartile associations with Lp(a) lipid concentrations for the African American (N=330), European American (N=381), and Hispanic-American (N=375) populations.

**B**. Allele 2 (shorter allele) CN quartile associations with Lp(a) lipid concentrations for the African American (N=330), European American (N=381), and Hispanic-American (N=375) populations.

**A**
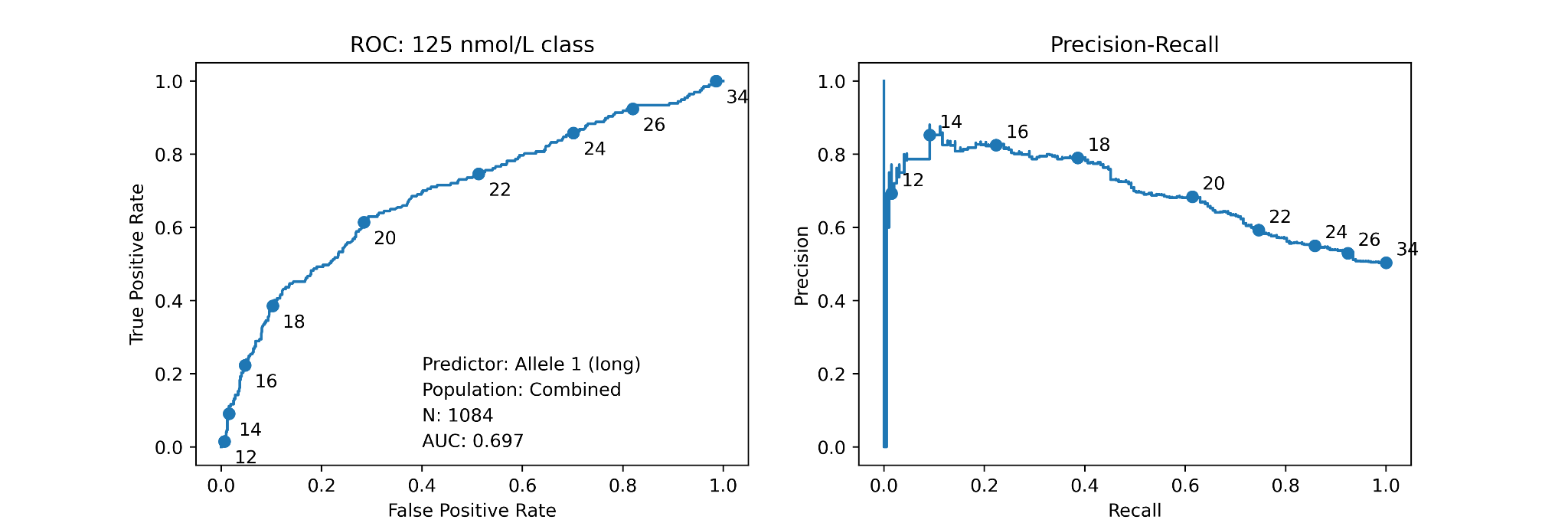

**B**
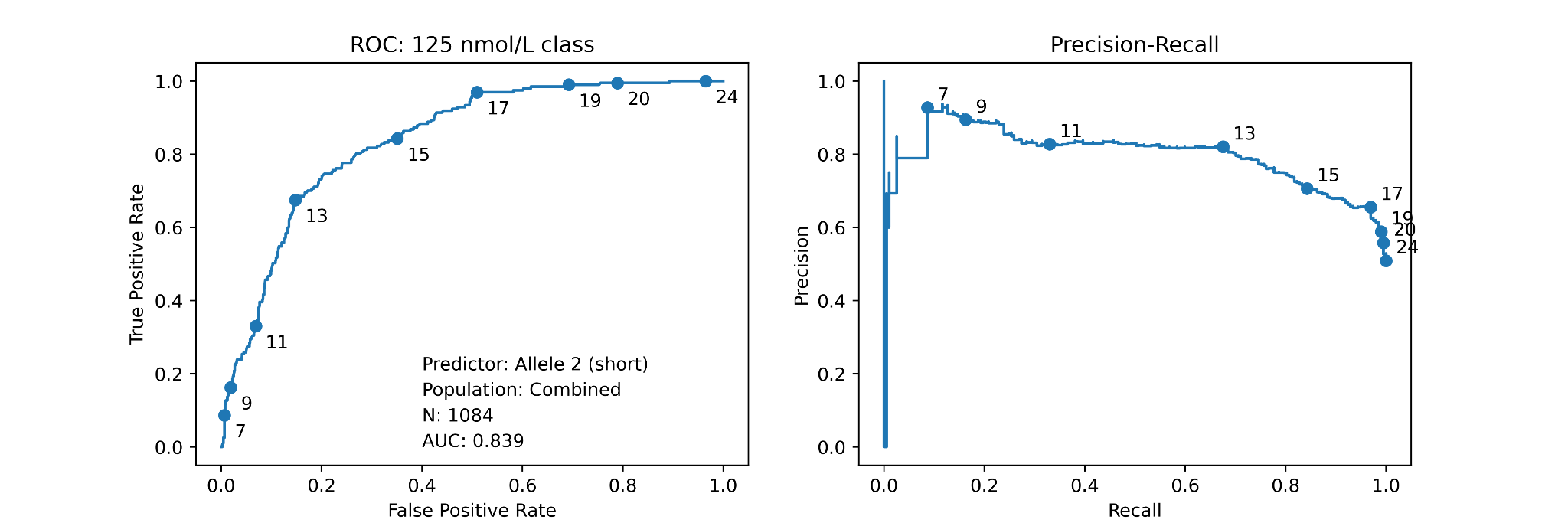

**C**
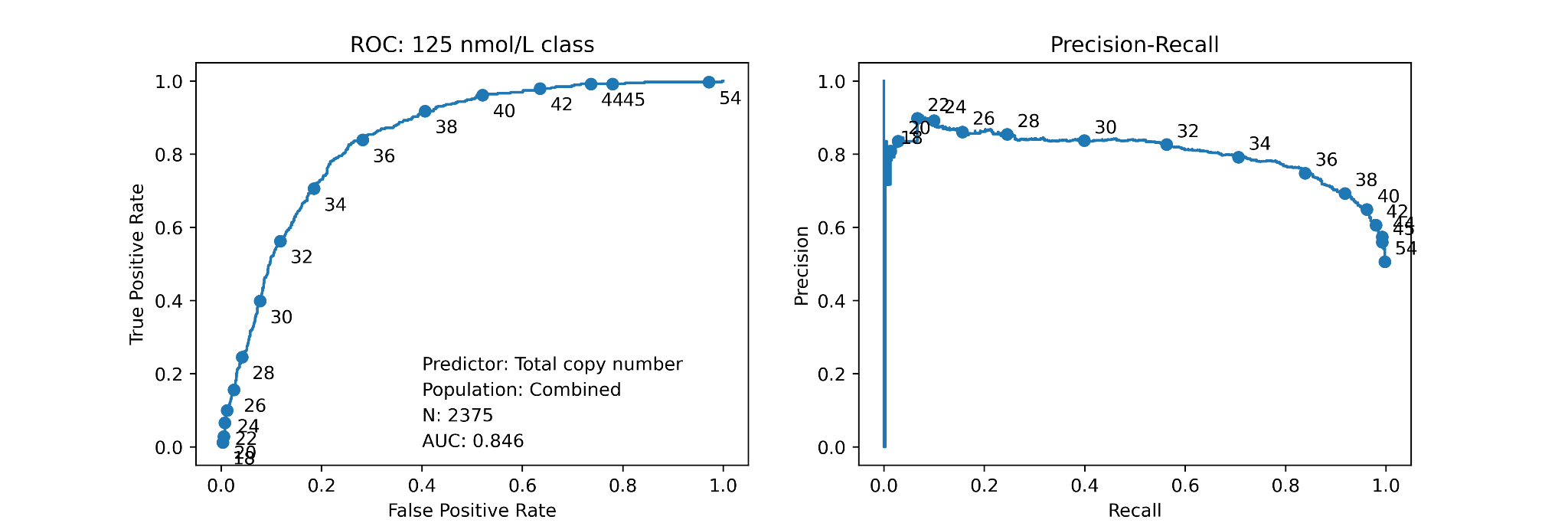

**Supplementary Figure S13: Analysis of KIV-2 copy number as predictor of high Lp(a).**

**A**. ROC Curve and Precision-Recall curve for classification of samples as high-Lp(a) (>125 nmol/L) using the Allele 1 CN.

**B**. ROC Curve and Precision-Recall curve for classification of samples as high-Lp(a) (>125 nmol/L) using the Allele 2 CN.

**C**. ROC Curve and Precision-Recall curve for classification of samples as high-Lp(a) (>125 nmol/L) using the total KIV-2 copy number.

Each plot contains samples from the African-American, European-American, and Hispanic-American populations.

**A**
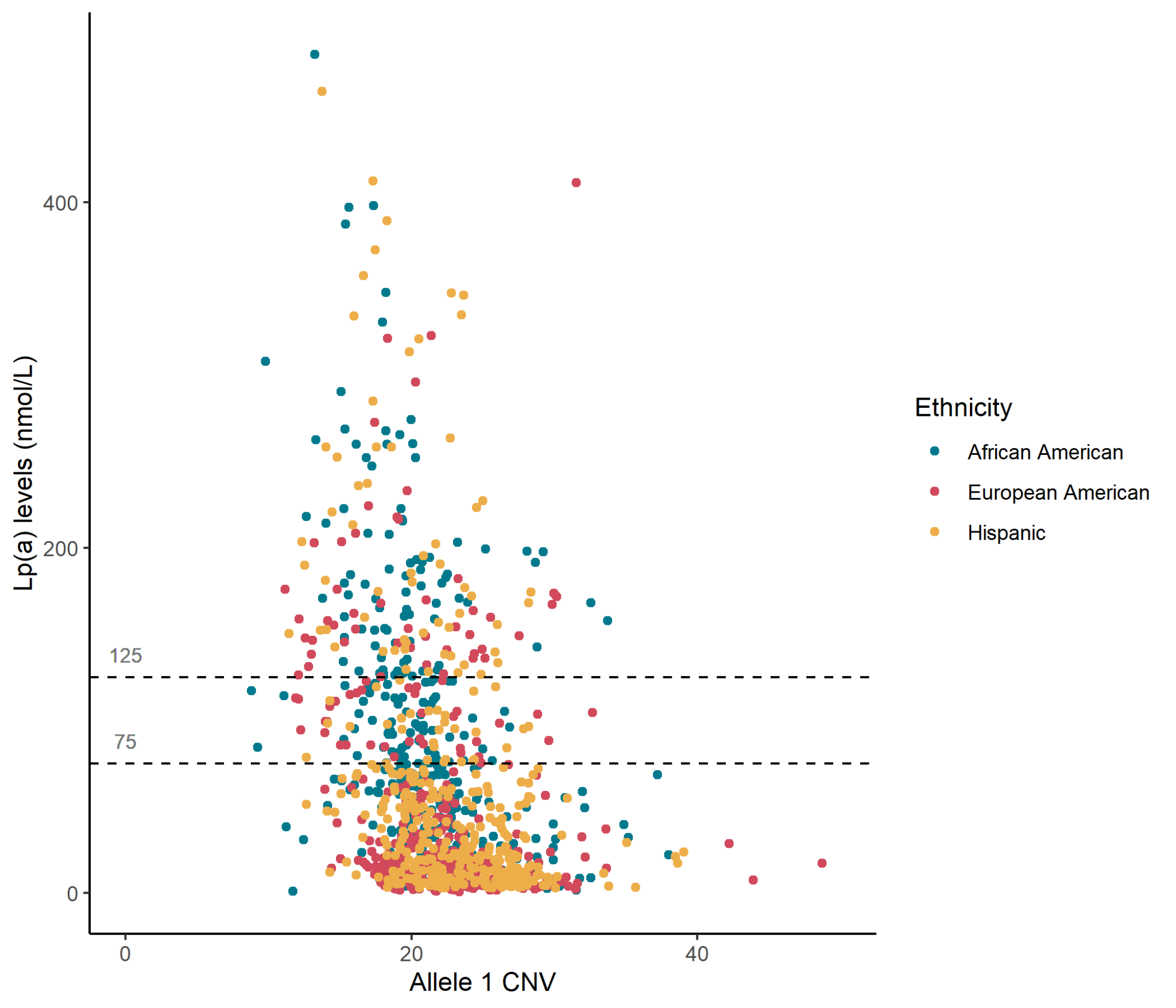

**B**

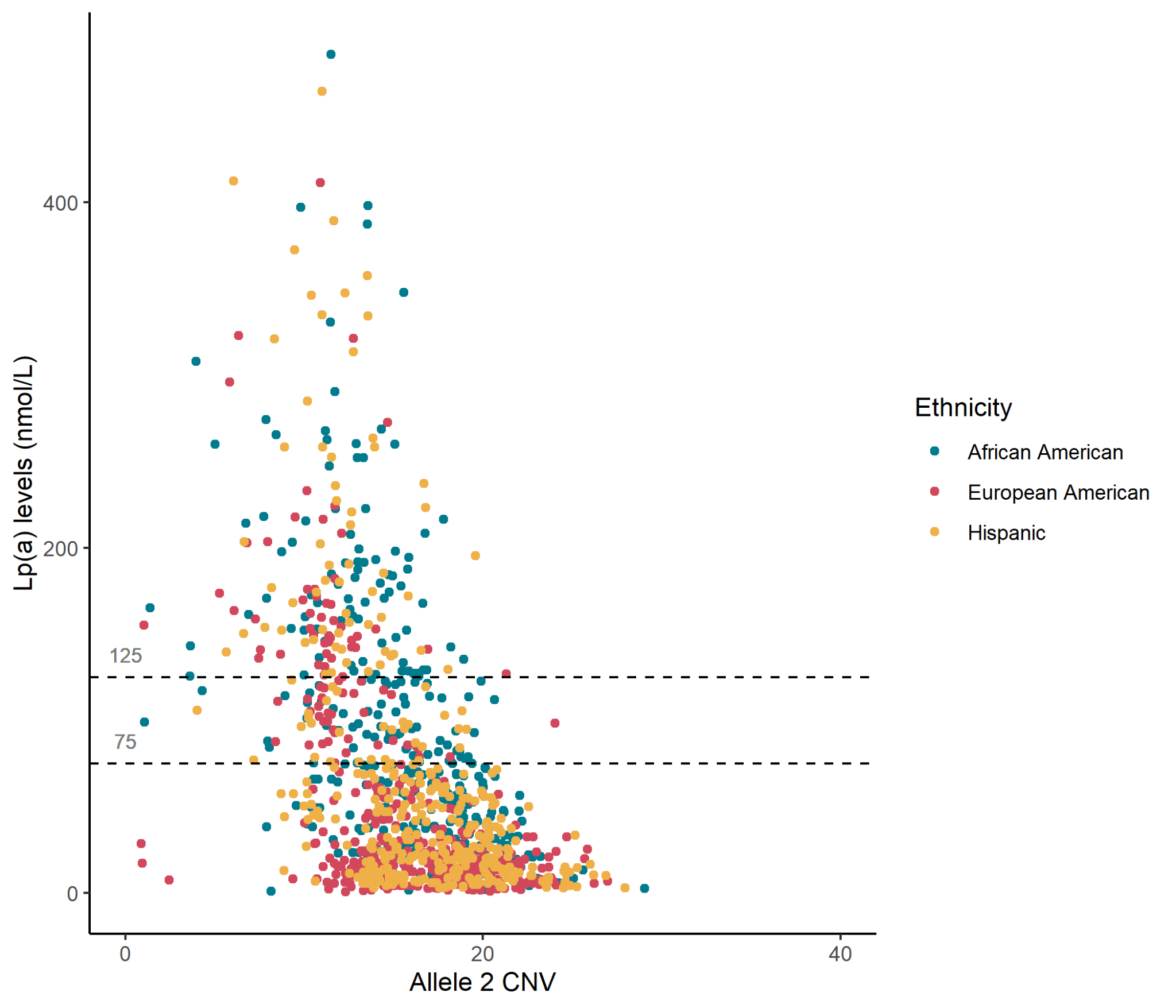

**Supplementary Figure S14: Allele length associations with Lp(a) concentrations**. **A**. Allele 1 (longer allele) CN and the corresponding Lp(a) concentrations. **B**. Allele 2 (shorter allele) CN and the corresponding Lp(a) concentrations
